# Impaired biogenesis of basic proteins impacts multiple hallmarks of the aging brain

**DOI:** 10.1101/2023.07.20.549210

**Authors:** Domenico Di Fraia, Antonio Marino, Jae Ho Lee, Erika Kelmer Sacramento, Mario Baumgart, Sara Bagnoli, Pedro Tomaz da Silva, Amit Kumar Sahu, Giacomo Siano, Max Tiessen, Eva Terzibasi-Tozzini, Julien Gagneur, Judith Frydman, Alessandro Cellerino, Alessandro Ori

## Abstract

Aging and neurodegeneration entail diverse cellular and molecular hallmarks. Here, we studied the effects of aging on the transcriptome, translatome, and multiple layers of the proteome in the brain of a short-lived killifish. We reveal that aging causes widespread reduction of proteins enriched in basic amino acids that is independent of mRNA regulation, and it is not due to impaired proteasome activity. Instead, we identify a cascade of events where aberrant translation pausing leads to reduced ribosome availability resulting in proteome remodeling independently of transcriptional regulation. Our research uncovers a vulnerable point in the aging brain’s biology – the biogenesis of basic DNA/RNA binding proteins. This vulnerability may represent a unifying principle that connects various aging hallmarks, encompassing genome integrity and the biosynthesis of macromolecules.

**One-Sentence Summary:** Translation pausing reshapes the aging brain proteome, revealing vulnerabilities in the biogenesis of nucleic-acid protein.

## Main text

Both aging and neurodegeneration disrupt protein homeostasis, also known as proteostasis, leading to the progressive accumulation of protein aggregates (*1*, *2*). Proteostasis involves mechanisms that regulate the coordination of protein synthesis, degradation, and localization and is essential to ensure an adequate supply of protein building blocks. It also prevents the accumulation of misfolded and "orphan" proteins susceptible to aggregation.

Age-dependent decline in proteostasis coincides with the onset of other aging hallmarks, but a causal connection remains unproven, necessitating integrative analysis to reveal these associations. This knowledge gap remains, at least partially, because these aspects were analyzed separately and in different model systems. Therefore, we conducted a comprehensive analysis of proteostasis in the aging brain of the short-lived killifish *Nothobranchius furzeri*. We chose killifish because of its accelerated aging and spontaneous emergence of neurodegenerative phenotypes (*3*–*8*)

Using this model system, we measured the effects of aging on the transcriptome, translatome, and multiple layers of the proteome, enabling quantification of their relationships. We also established a protocol for long-term partial inhibition of proteasome activity to investigate which age-related brain phenotypes are caused by this specific dysfunction *in vivo*. Finally, we performed Ribo-Seq to assess the contribution of mRNA translation to proteome alterations. Our work reveals that the biosynthesis of specific proteins rich in basic amino acids becomes perturbed with age due to aberrant translation elongation and pausing. Such alterations lead to the depletion of protein complexes involved in DNA/RNA-binding and protein synthesis, and remodel the proteomes of organelles such as mitochondria. This altered translation dynamic also provides a mechanistic explanation for the highly conserved age-related discrepancies between transcriptome and proteome changes (*9*,*10*,*11*,*12*,*13*,*14*,*6*), which have been linked to neurodegeneration in humans (*15*). Thus, our work reveals how the biogenesis of a specific subset of proteins might further enhance vulnerabilities of the proteostasis system in the aging brain and contribute to the exacerbation of other aging hallmarks.

### Loss of basic proteins in the aging brain

To investigate the disruption of protein homeostasis in aging, we focused on the loss of correlation between changes in gene transcripts (mRNA) and corresponding protein, an intra-species conserved phenomenon (*9*,*10*,*11*,*12*,*13*,*14*,*6*) here referred to as "decoupling", for which a biological explanation is still lacking. To address the origin of decoupling, we combined the quantification of age-related changes in gene transcripts and protein levels obtained from RNA sequencing (RNAseq) and mass spectrometry-based proteomics data, respectively (Figure 1A-B, Figure S1A-H). We calculated a "decoupling score" by measuring the differences between protein and transcript changes (see methods). The decoupling score effectively describes discrepancies between transcript and protein changes by identifying subsets of proteins displaying “positive protein-transcript decoupling”, i.e., protein level higher than expected from changes of its corresponding transcript, or “negative protein-transcript decoupling”, i.e., protein level lower than expected from changes of its corresponding transcript (Figure 1B-C, Table S1).

**Figure 1:**
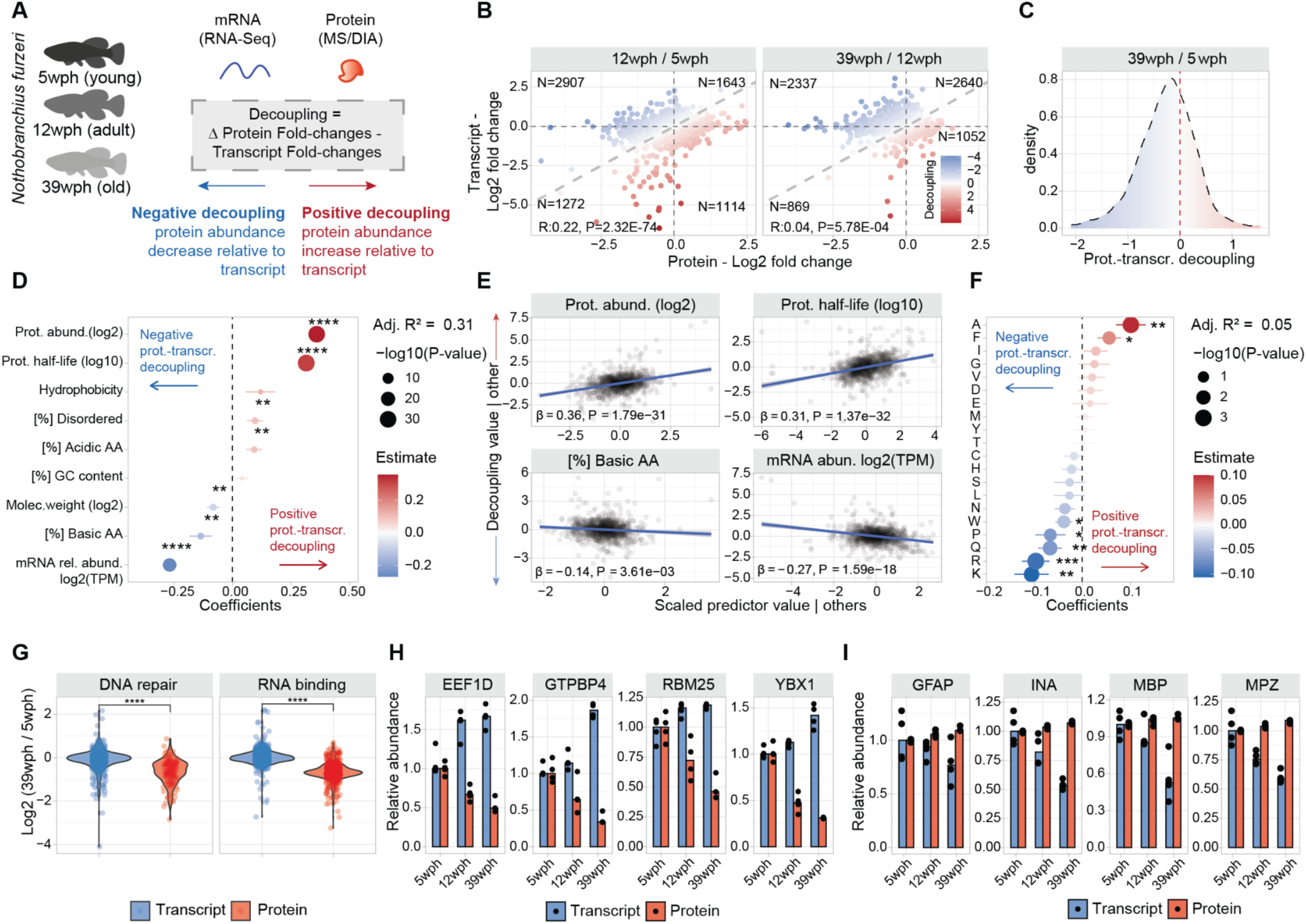
Protein-transcript decoupling affects highly abundant and basic proteins in opposite manners. A) Characterization of protein-transcript decoupling in the aging brain of killifish. Positive decoupling values indicate increased protein abundance relative to transcripts, while negative decoupling indicates decreased protein abundance compared to transcripts. B) Scatterplot comparing protein and transcript fold changes in aging brain. Color represents decoupling score, red – increased protein abundance relative to transcript, blue – decreased protein abundance compared to transcript. Grey dashed lines indicate equal changes. C) Density distribution of decoupling scores for 39 wph vs. 5 wph comparison. Red: positive decoupling, Blue: negative decoupling. D) Multiple linear regression analysis of decoupling scores based on transcript and protein features. asterisks represent the -log10 P-values of the F-test. E) Added variable plot between features and decoupling scores. F) Multiple linear regression analysis of decoupling scores based on protein amino acid composition. asterisks represent the -log10 P-values of the F-test. G) Transcript and protein fold changes for RNA binding and DNA repair proteins. Two-sample Wilcoxon test H and I) Examples of proteins with negative (H) and positive (I) decoupling (N=3-4). *P ≤ 0.05; **P ≤ 0.01, ***P ≤ 0.001, ****P ≤ 0.0001. Related to Figure S1 and Table S1.

The obtained scores displayed a median shift towards negative values (Figure 1C) due to an overall skew towards negative fold changes at the proteome level (Figure S1D), which was independent of sample normalization (Figure S1C). To assess the reproducibility of this metric, we compared the decoupling scores from this study to the ones obtained using an independent brain aging dataset that we previously generated (*6*). Supporting our observations, there was a significant positive correlation between these datasets (Figure S1I), despite technical differences in the quantitative proteomics workflows: tandem-mass tags (TMT) based quantification (*6*) compared to label-free Data Independent Acquisition (DIA, this study).

We then applied a multiple linear regression model to interrogate the association between the measured decoupling scores (response variable, N=1188 complete observation) and distinct biophysical properties of transcripts and proteins (N=9 features). Our model explained 31% of the decoupling variance (Adjusted *R^2^* = 0.31, Figure 1D). We detected estimated protein absolute abundance (see methods, **β**=0.36, *P* < 2.20E-16) and protein half-life (as described in (*16*), **β**=0.31, *P* < 2.20E-16) as the parameters with the highest correlation with higher protein levels than expected from transcript changes (Figure 1E). On the other hand, the parameters with the highest correlation with negative decoupling (i.e., lower protein levels than expected from transcript changes) were relative transcript abundance (expressed as log2 transcripts per million (TPM) **β**=-0.26, *P* < 2.20E-16) and proportion of basic amino acids (**β**=-0.13, *P* = 4.30E-03, Figure 1D-E). To understand the contribution of amino acid composition to the observed discrepancies, we employed a second regression model with protein amino acid composition as the sole predictor variable. Our analysis revealed significant correlations between negative decoupling and the content of lysine, proline, glutamine, and arginine (Figure 1F).

Intrigued by these findings, we investigated the behavior of proteins involved in DNA repair and RNA-binding, due to their high content of basic amino acids. Both these groups of proteins showed an age-dependent decrease of protein- but not transcript levels (Figure 1G and H). On the other hand, myelin components, e.g., myelin basic protein (MBP) and myelin protein P0 (MPZ), and intermediate filament proteins, e.g. glial fibrillary acidic protein (GFAP) and alpha-internecine (INA), showed decreased transcript- but not protein-levels with aging (Figure 1I), consistently with their long half-lives and low turnover rates. These results show that specific classes of proteins experience discrepancies between protein and transcript levels in the aging vertebrate brain. The distinct biophysical and biochemical characteristics promoting these discrepancies suggest the presence of shared molecular attributes that might drive these phenomena. In particular, our data highlight a potential mechanistic link between the content of basic amino acids and the age-related decrease in protein levels compared to the corresponding mRNAs.

### Convergence of proteome alterations on ribosomes and respiratory chain complexes

We then explored how discrepancies between transcript and protein changes relate to other proteome alterations. To do so, we employed a comprehensive approach characterizing age-related changes in proteome insolubility by implementing a differential detergent extraction protocol (Figure 2A, Figure S2 and Table S2, see methods (*17*)) and organelle composition, using subcellular fractionation in combination with mass-spectrometry (*18*) (Figure S3 and Table S2, see methods). We additionally quantified changes in phosphorylation, ubiquitylation, and acetylation to detect how the landscape of post-translational modifications is affected in the aging brain (Figure 2A, Figure S4 and Table S3, see methods). The combined data showed changes in organelle proteome composition, proteasome solubility, alterations in the post-translational modification in the aging brain (Figure S5, see Supplementary text), and age-related proteome alterations in proteins encoded by genes genetically linked to neurodegeneration in humans (Table S4, see Supplementary text and Figure S6).

**Figure 2:**
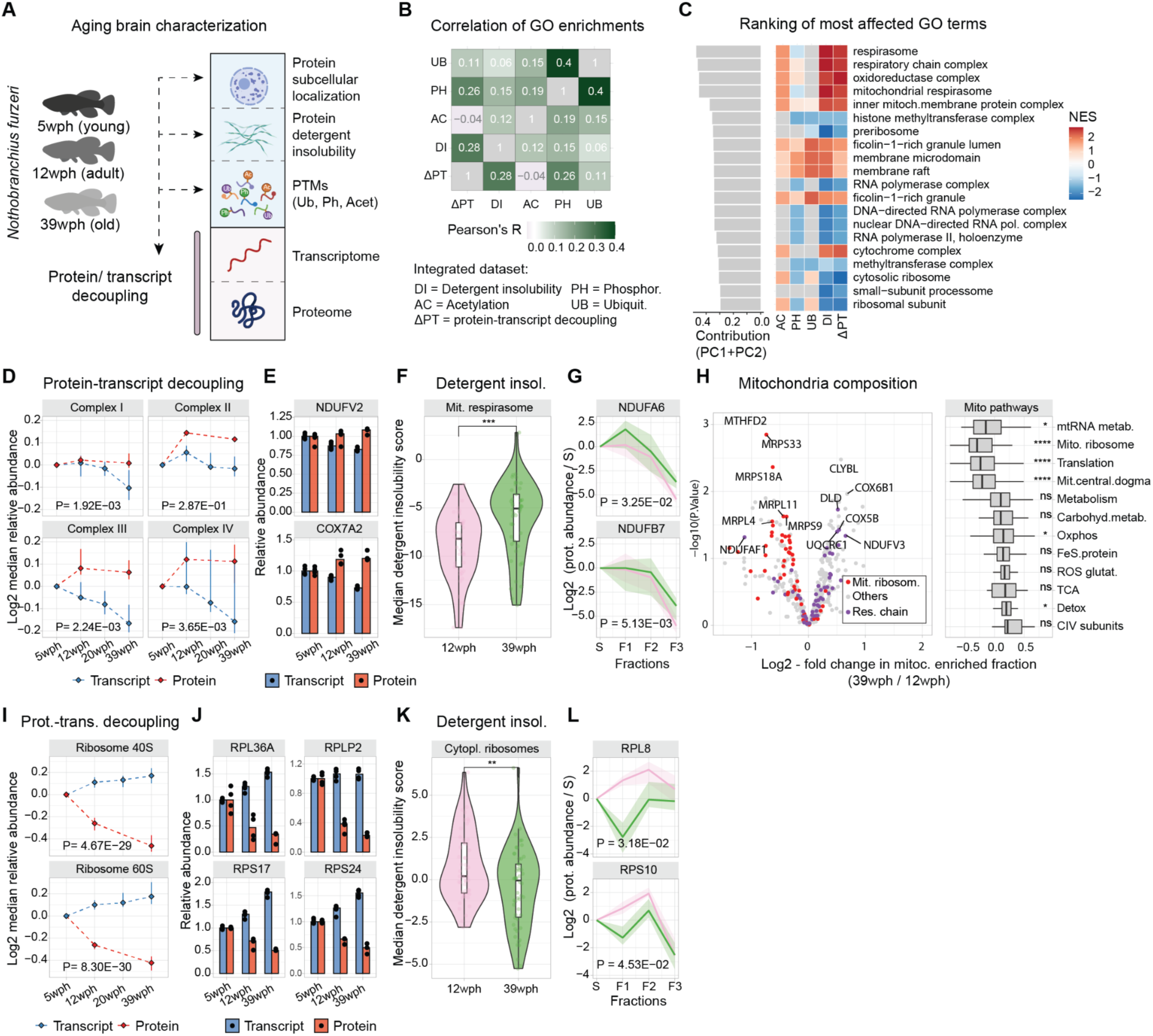
Proteome alterations converges on ribosomes and respiratory chain complexes. A) Overview of the datasets generated at the beginning of this study (wph= weeks post-hatching). B) Heatmap shows correlations of normalized enrichment scores (NES) across datasets (DR=Detergent insolubility, ΔPT=protein-transcript decoupling, AC=Acetylation, PH=Phosphorylation, UB=Ubiquitylation). C) Top-ranking GO terms with strong contributions to PCA analysis. D) Line plots for respiratory chain proteins’ transcript (blue) and protein (red) median abundance across age groups (N=3-4, MANOVA). E) Examples of respiratory chain proteins with positive protein-transcript decoupling. F) Violin plot for detergent insolubility scores of mitochondrial respirasome proteins (N=4, two-sample Wilcoxon test). G) Examples of detergent insolubility profiles for respiratory chain proteins with increased detergent insolubility during aging (N=4, MANOVA). H) Volcano plot for changes in mitochondrial proteome due to aging. Box plot shows the effect of aging on different groups of mitochondrial pathways (N=4, two-sample Wilcoxon test). I) Ribosomal proteins’ transcript and protein abundance across age groups (N=3-4, MANOVA). J) Examples of ribosomal proteins displaying negative protein-transcript decoupling (N=3-4). K) Violin plot for detergent insolubility scores of cytoplasmic ribosomal subunits (N=4, two-sample Wilcoxon test). L) Examples of detergent insolubility profiles for ribosomal proteins with decreased detergent insolubility during aging (N=4, MANOVA). *P ≤ 0.05; **P ≤ 0.01, ***P ≤ 0.001, ****P ≤ 0.0001. Related to Figure S2:S7 and Tables S2,S3.

To explore the relationship among different types of proteome alteration in the aging brain, we conducted a gene set enrichment analysis on the age-related proteome changes for each of the generated datasets (see methods). By calculating Pearson’s correlation coefficient between enrichment scores across datasets, we found a positive correlation between protein-transcript decoupling and increased detergent insolubility, a hallmark of protein aggregation (Pearson’s R = 0.28, P < 2.20E-16), as well as protein phosphorylation (Pearson’s R = 0.26, P = 6.67E-08), while other alterations, for instance, changes in protein ubiquitylation, showed a smaller correlation value (Pearson’s R = 0.11, P = 1.23E-02, Figure 2B).

To unbiasedly identify the most prominently affected cellular components in our analysis, we then used principal component analysis (PCA) to summarize the normalized enrichment scores (NES, see methods) and ranked GO terms by calculating the values of their projections on the first two principal component axes. We found that the highest-ranking terms were related to components of the mitochondrial respiratory chain and ribosomes (Figure 2C). These two sets of protein complexes were often affected by aging in opposite ways (Figure 2C). Components of the respiratory chain showed a progressive decrease in their transcripts together with a stable or modest increase of the corresponding protein levels (Figure 2D-E, Figure S7B). Respiratory chain proteins also showed an overall increase in detergent insolubility with aging (Figure 2F-G). Importantly, these alterations primarily affected respiratory chain components but not mitochondrial proteins in general (Figure S7C). To corroborate these findings, we interrogated our subcellular fractionation data (Figure S3). This analysis allowed us to identify two key aspects: (i) changes in the protein composition of aged mitochondria, notably a significant decrease in the relative abundance of mitochondrial ribosomes and an increase in the relative abundance of oxidative phosphorylation (Figure 2H and Figure S7D), and (ii) altered subcellular distribution of specific mitochondrial proteins (Figure S7E-F). These analyses provide support for a global remodeling of the mitochondria during aging.

In contrast, both cytosolic and mitochondrial ribosomal protein levels progressively decreased during aging (reaching, on average, a ∼25% reduction in old brains), while their corresponding transcripts increased (Figure 2I-J, Figure S7G-H). The reduced level of ribosomal proteins was accompanied by a decreased detergent insolubility (Figure 2K-L, Figure S7H-I). This alteration might be related to the loss of ribosome stoichiometry and partial mis-/disassembly that we previously described in the old killifish brain (*6*). Interestingly, we noticed similar patterns for other large complexes rich in basic amino acids, like RNA polymerase II (Figure S7J-K), that might indicate common mechanisms altering the homeostasis of these key complexes. These combined analyses reveal that the mitochondrial respiratory chain, ribosomes, and other protein complexes enriched in basic amino acids are preferentially and differently affected in the aging brain. A web-based application is available for the exploration of these datasets (https://genome.leibniz-fli.de/shiny/orilab/notho-brain-atlas/ credentials username: reviewer password: nothobrain2023)

### Impact of proteasome impairment on the brain proteome

Protein degradation by the ubiquitin-proteasome system regulates protein levels in organelles and complexes, including ribosomes and mitochondria. Previous studies (*6*,*2*,*19*) have linked aging with a decline in proteasome activity. To study the impact of proteasome activity on the aging brain, we simulated its impairment by chronically reducing its activity in adult killifish. We optimized *in vivo* dosage of bortezomib, a dipeptide that binds with high affinity and blocks the catalytic site of the proteasome, to maintain a ∼50% inhibition in the brain of adult killifish over 4 weeks without inducing overt toxicity (Figure 3A, Table S5). GO enrichment analysis revealed brain proteome and transcriptome adaptive responses to proteasome inhibition, including over-representation of proteasome-related terms (Figure 3A) and specific proteostasis network alterations (Figure S8A). These include increased protein levels of proteasome activators (PSME1/PA28ɑ, PSME3/PA28γ, and PSME4/PA200) and increased mRNA levels of key autophagy genes such as *ATG5* and *ATG7* (Figure 3B). Some responses, like elevated HSPB1 and HSPA6, arose spontaneously in aged killifish brains (Figure 3B). Immunofluorescence analysis of lysosomes revealed a marked increase in their area, volume, and sphericity (Figure 3C), a phenotype typical of aging (Figure S8B), lysosomal storage disorders (*20*), and neurodegenerative diseases (*21*). Proteasome inhibition globally reduced mitochondrial protein levels independent of transcription (Figure 3D) and without altering master regulators of mitochondrial genes (Figure S8C). Similar to aging it also induced a reduction of mitochondrial content (estimated calculating ratio of mitochondrial DNA (mtDNA) to nuclear DNA, Figure 3D-S8D).

**Figure 3:**
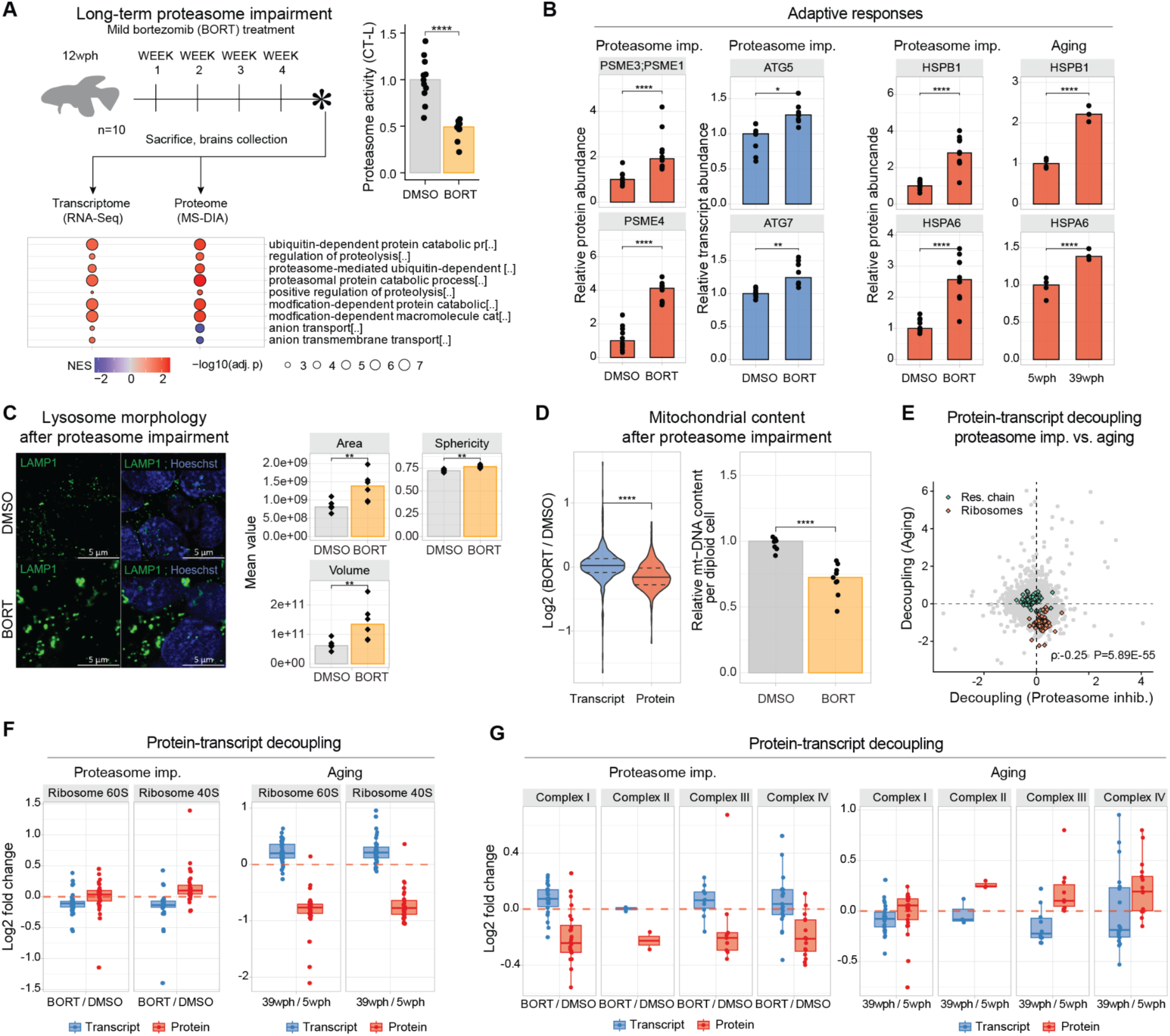
Effects of four weeks *in vivo* proteasome impairment on the adult killifish brain. A) Adult killifish (12 wph, N=10) received weekly intraperitoneal injections of proteasome inhibitor bortezomib or DMSO control. Bottom panel: Gene Set Enrichment Analysis (GSEA) color-coded by normalized enrichment score (NES). Top-right: Chymotrypsin-like proteasome activity quantification (two-sample Wilcoxon test, N=10). B) Barplot: Protein (red) transcript (blue) quantity in proteostasis network proteins in aging and proteasome inhibition (two-sample T-test, N=10). Asterisks indicate Q.value (protein) and Adjusted P.Value (transcript). C) Left: Lysosome (LAMP1) immunofluorescence. Scale bars = 5μm; right: Lysosome morphology analysis (two-sample T-test N=6). D) Left: Proteasome effect on mitochondrial transcripts and proteins (two-sample Wilcoxon test); right: Relative mtDNA copy number was calculated using real-time quantitative PCR with primers for 16S rRNA mitochondrial gene and Cdkn2a/b nuclear gene for normalization (N=10, two-sample Wilcoxon tests). E) Decoupling scores comparison between aging and proteasome impairment for respiratory chain (green) and ribosomal (orange) proteins (Spearman correlation was selected due to the presence of outliers in the distribution). F) Ribosome decoupling comparison between aging and proteasome impairment. G) Oxidative phosphorylation protein decoupling comparison between aging and proteasome impairment. *P≤0.05; **P≤0.01, ***P≤0.001, ****P≤0.0001. Related to Figure S8, Table S5.

As expected, proteasome impairment led to an increased abundance of shorter-lived proteins, consistent with its role in regulating protein turnover (*22*)(Figure S8E). We then combined RNAseq and proteome data to understand the contribution of reduced proteasome activity to the age-related discrepancies between transcript and protein levels. We showed that proteasome impairment leads to decoupling between transcript and protein changes (Figure S8F, Table S5). However, when we applied the same linear regression models used for aging, we found distinct biophysical properties associated with decoupling by bortezomib compared to aging (Figure S8F). Specifically, proteasome inhibition caused an accumulation of basic proteins, independently of transcription (Figure S8F), and decoupling scores induced by proteasome impairment and aging were surprisingly negatively correlated (Spearman Rho = -0.25, P < 2.20E-16, Figure 3E). This included the cases of ribosomes (Figure 3F) and respiratory chain complexes (Figure 3G). Thus, our data reveal specific alterations induced in the adult killifish brain by partial proteasome impairment, some of which recapitulate aging brain phenotypes. Nonetheless, reduction in proteasome activity does not appear to be directly responsible for the loss of basic proteins observed in the old brains, hinting at other possible mechanisms.

### Aberrant translation pausing correlates with decreased levels of basic proteins in old brains

Our findings show that imbalances in proteostasis during aging go beyond proteasome dysfunction. Other factors, such as differential mRNA translation at old age, could cause the observed discrepancies between transcript and protein levels. Therefore, we explored this hypothesis with a Ribo-Seq experiment in aging killifish brains (Figure 4A, Table S6). Tri-nucleotide periodicity and consistent replicates (Figure S9A-B) showed the overall quality of the data, while comparison between mRNA levels and ribosome occupancy showed the expected correlation between the two (R=0.25, *P* < 2.20E-16, Figure S9C). When we estimated translation efficiency (TE, see methods), we observed that this measure was better at explaining protein changes (R=0.32, P < 2.20E-16, Fig. 4B) in comparison to transcript changes (R=0.23, P < 2.20E-16), consistent with previous observations in mammals (*23*). For instance, shifts in TE led to consistent changes in protein levels for certain protein complexes, such as the Complex IV of the respiratory chain, and the 26S proteasome (Figure 4C-D and S9D). Interestingly though, ribosomes, RNA polymerase II, and other nucleic-acid binding proteins linked to DNA repair (Figure 4C-D, and S9D) didn’t exhibit the same behavior, excluding TE as the cause for their decreased abundance.

**Figure 4:**
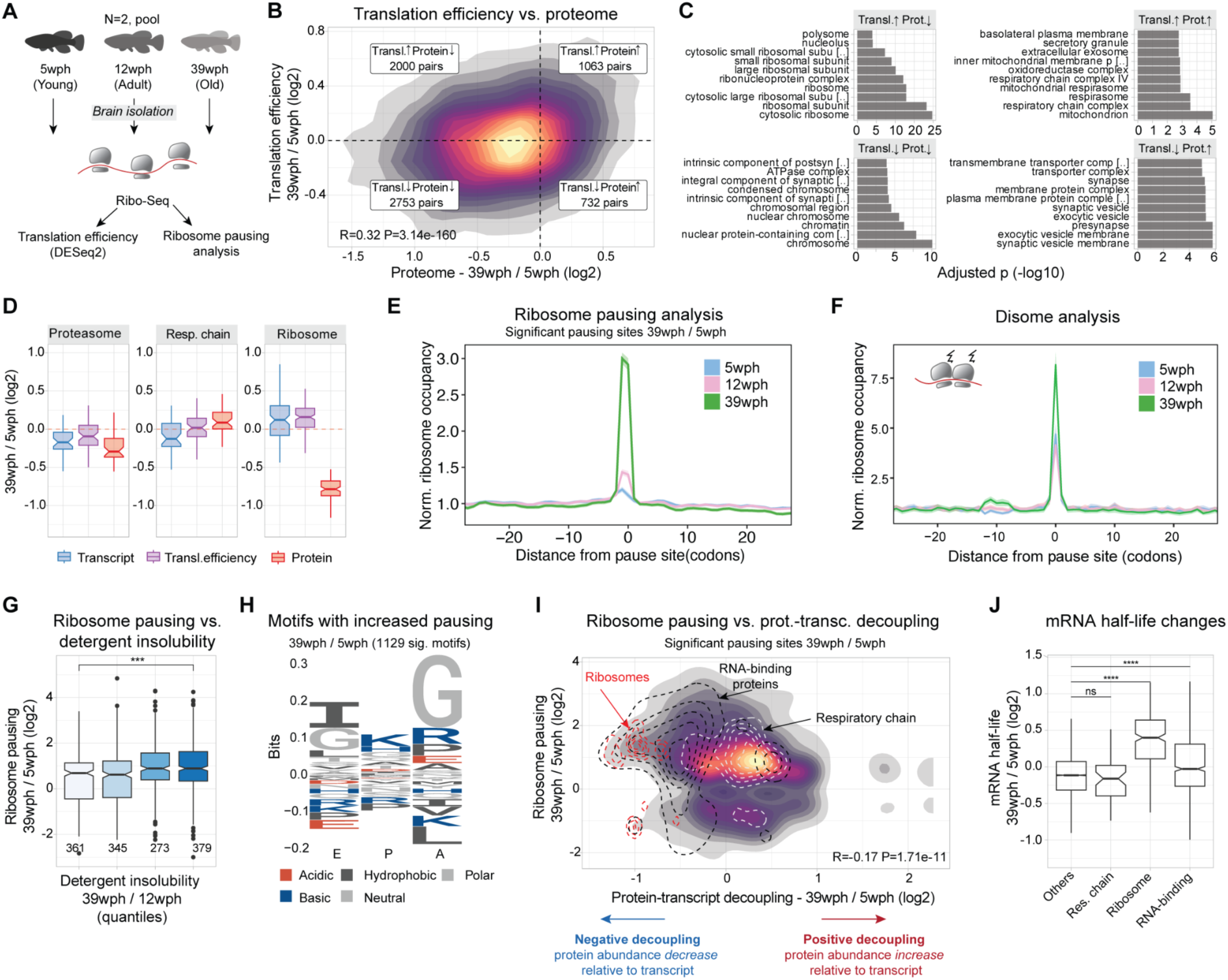
Increased translation pausing in the aging killifish brain. A) Experimental Workflow: Ribosome profiling was conducted on brains of *Nothobranchius furzeri* at different ages—Young (5 weeks post-hatch, wph), Adult (12wph), and Old (39wph). Each age group had two replicates, each consisting of pooled samples from 10-15 animals. B) A 2-D density plot illustrates the connection between age-induced changes in protein abundance (x-axis) and alterations in translation efficiency (y-axis). Different quadrants highlight modes of translation regulation. C) GOEnrichment analysis (ORA) for each quadrant from B. x-axis: -log10(adjusted P-value) Fisher test, Holm correction. D) Differential regulations for key complexes, 26S Proteasome, oxidative phosphorylation, and cytoplasmic ribosomes – Transcriptome (blue), Translation efficiency (purple), and Proteome (red), in Old vs. Young. E) Lineplot showing the normalized ribosome distribution at pausing sites across different age groups. F) Lineplot depicts normalized disome ribosome distribution at disome pausing sites for various age groups. G) Boxplot showing solubility vs. ribosome pausing. x-axis: solubility quantiles (25% of total distribution each), y-axis: log2 fold changes in pausing for significant sites (Adj. P-value < 0.05). Numbers indicate observations. Two-sample Wilcoxon tests H) Peptide motif associated with age-dependent increased pausing (Pause score at 39wph > Pause score at 5wph and 12wph, and Pause score at 39wph > 6). y-axis: relative residue frequencies, x-axis: ribosome positions (E, P, A). I) 2-D density plot showing relation between significant pausing changes (Adj. P-value < 0.05) on y-axis and decoupling metrics (x-axis). Contours: cytoplasmic ribosomes (red), RNA-binding proteins (black), oxidative phosphorylation (white). J) Boxplot showing mRNA half-life estimate changes (methods) between 39 wph and 5 wph. x-axis: selected categories. Asterisks: two-sample Wilcoxon test, Holm correction. *P ≤ 0.05; **P ≤ 0.01, ***P ≤ 0.001, ****P ≤ 0.0001. Related to Figure S9, Table S6.

To investigate further this intriguing aspect, we drew inspiration from studies on aged nematodes and yeasts, which exhibit age-related impaired translation elongation and ribosome pausing (*24*). We queried our Ribo-Seq data for signatures of translation pausing (see methods) and revealed an overall increase in site-specific pausing in the aging brain (Figure 4E, Table S6), with disome analysis confirming increased ribosome collisions (Figure 4F). Anisomycin-induced ribosome stalling in killifish cells (Figure S9E) showed a higher molecular weight ubiquitylated band in the immunoblot against the 40S subunit RPS3, commonly associated with ribosome collision (*25*–*27*). Aged brains showed similar changes (Figure S9F), even though most ribosomal protein ubiquitination decreased with age (Figure S9G). Similarly to aged yeast and nematodes, we also observed a decrease in a subset of proteins involved in ribosome quality control (RQC) (Figure S9H). This RQC reduction may worsen ribosome collisions and stalling as age progresses, potentially slowing stalled mRNA degradation and causing their accumulation in aging cells. When we investigated the impact of translation pausing on cellular proteostasis, we reported a clear association between translation pausing and increased detergent insolubility (Figure 4G). More interestingly, these alterations affected key proteostasis network components such as the proteasome (Figure S9I), hinting at a possible vicious cycle of proteostasis collapse. Stretches enriched in codons for basic residues (arginine and lysine), as well as glycine, were enriched at both pausing (Figure 5H) and disome sites (Figure S9J). Notably, our decoupling model linked the same residues (arginine and lysine) to reduced protein levels (Figure 1F). Furthermore, we found a significant correlation between pausing and protein-transcript decoupling (Figure 4I, R=-0.17, P < 2.20E-16), explaining cases for ribosomal and RNA-binding proteins, where protein decline does not follow transcript changes. On the contrary, components of the respiratory chain did not show any remarkable deviation from the overall pausing distribution (Figure 4I), although different complexes showed distinct pausing profiles (Figure S9K).

**Figure 5:**
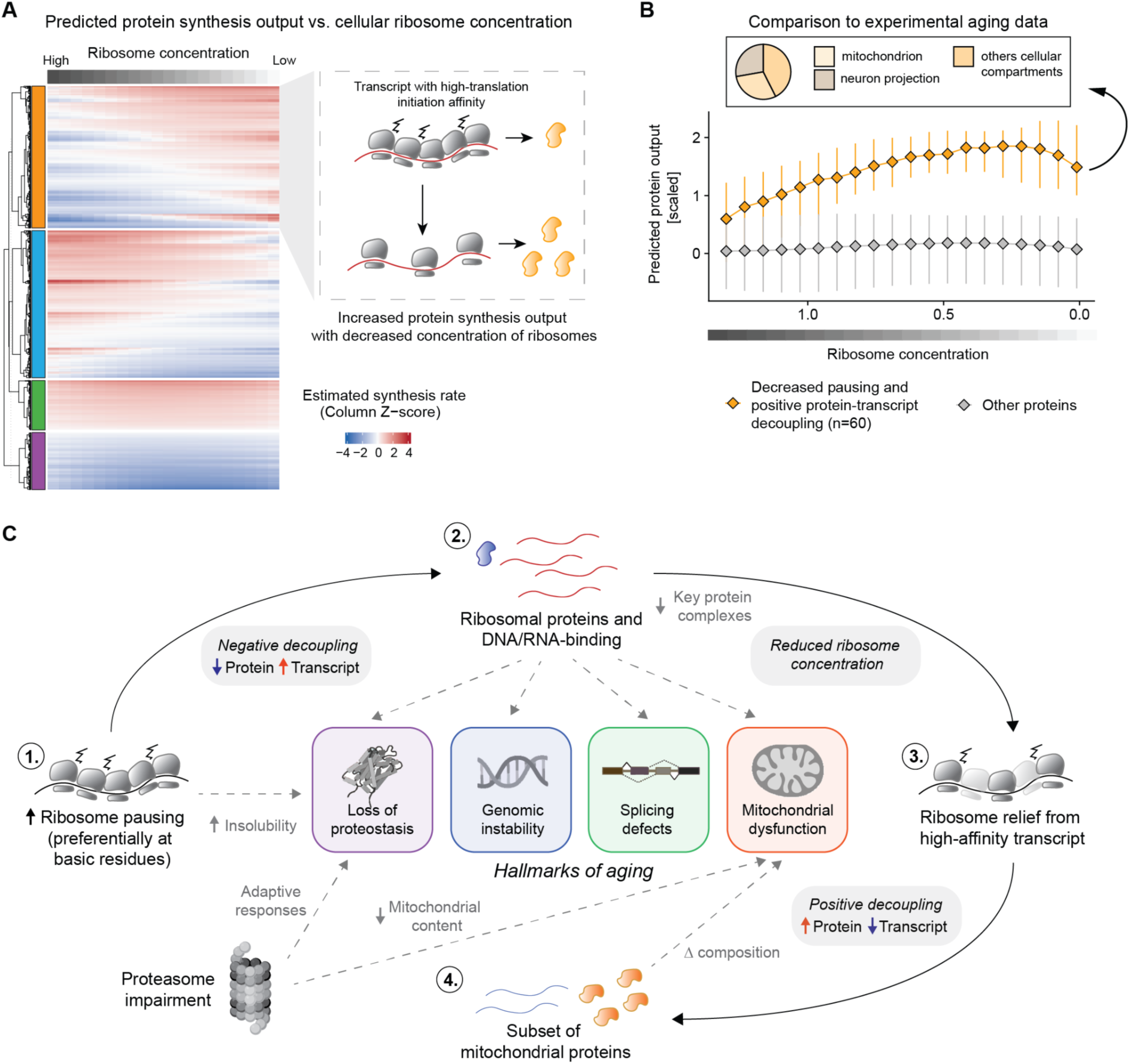
Reduced ribosome levels can lead to translation reprogramming in the aging brain. A) Heatmap showing the estimated protein output, modeled as described in Mills and Green 2017 (*32*). Each column in the heatmap indicates the estimated protein output for a specific ribosome concentration. Transcripts are clustered with a hierarchical clustering using the “ward D2” algorithm on the dissimilarity (1 - Person’s correlation) measure. For display purposes, the heatmap represents 5000 rows randomly sampled from all datasets. In the right panel, an illustrative example of a cluster displaying increased estimated protein output as a function of reduced ribosome levels. For these transcripts, the general ribosome decrease is predicted to relieve trafficking and pausing, leading to overall improved protein production. B) Lineplot showing the estimated protein output for transcript displaying decreased ribosome pausing in the Ribo-Seq data (median per transcript log2 Pausing 39 wph / 5 wph < 0 and Adjusted P-value <=0.15) and increased protein levels relative to the transcript in the decoupling model (orange). The x-axis represents the simulated decreased ribosomal concentration, while the y-axis indicates the estimated protein output, as shown also in A. C) Schematic representation of the translation reprogramming model and its connection with the relevant hallmarks of aging. Aging is associated with increased ribosome collision and pausing on ribosomal proteins, leading to a ∼25% reduction of ribosome levels. This generalized decrease of available ribosomes could drive the translation of other high-affinity mRNAs leading to increased protein levels in the aging brain. Related to Table S7.

Changes in translation and pausing affect mRNA half-life (*28*–*30*). We additionally investigated the association between translation pausing and mRNA half-life, by estimating it from RNA-Seq data (see methods, (*31*)). This led to the discovery that, in old brains, transcripts encoding for ribosomal proteins and RNA-binding proteins show an increased half-life compared to the rest of the transcriptome (Figure 4J). This highlights that alteration in translation might influence other general changes in the transcriptome’s structure. In summary, our results show that increased ribosome occupancy does not necessarily result in enhanced protein synthesis in the aging brain, of note we propose that translation dysfunction may represent the underlying cause for the decreased levels of ribosomal proteins and other nucleic-acid binding proteins (enriched in basic amino acids) in the aging brain. Such reduction might further exacerbate other aging hallmarks that depend on the activity of these proteins.

### A possible model for protein biosynthesis in the aging brain

The results presented so far point to alterations in protein synthesis in old brains leading to a reduction of ribosomal proteins, among others. We hypothesized that the ensuing lower levels of ribosomes, particularly in light of increased load on the RQC machineries, may in turn lead to a vicious cycle of dysfunction. Altered ribosome concentration has been known to directly impact the translation of specific mRNAs, as observed in a group of inherited diseases collectively referred to as ‘ribosomopathies’ (*32*, *33*). We thus attempted to extend a model proposed by (*32*, *33*) to the aging scenario. The original model predicts that the protein output of specific mRNAs can be influenced by ribosome availability depending on transcript-specific translation initiation rate *k*_*i*_ (where *k*_*i*_ refers to the affinity of specific mRNAs sequences to bind ribosomes) (*32*, *33*). Under these assumptions, a decrease in ribosome concentration can, for example, increase protein synthesis from transcripts that have a high translation initiation rate by lowering the total ribosome load on them and therefore relieving trafficking and pausing events (Figure 5A). To test this hypothesis in the context of aging brain, we estimated *k*_*i*_ from killifish 5’-UTR sequences based on experimental data (*34*), and modeled the estimated synthesis rate as described in (*32*, *33*) (see methods, Figure 5A, Table S7). In agreement with the model, a subset of killifish transcripts displayed an increase in predicted synthesis rate as a function of decreased ribosome concentration (orange cluster in Figure 5A and Table S7). To test these predictions on our experimental data, we selected a specific set of proteins showing decreased translation pausing and increased protein abundance in our decoupling model (60 proteins, bottom right quadrant in Figure 4I). We then estimated their predicted synthesis rates as a function of ribosome concentration. Consistent with the experimental data, the relative synthesis of this subset of proteins was predicted to increase following a reduction of ribosome concentration (Figure 5B). Approximately one-third of these proteins were mitochondrial (including 7 components of the respiratory chain), and another prominent fraction belonged to proteins related to neuron projections (Figure 5B). Intriguingly, the absence of ribosomal proteins in this subset, despite their high *k*_*i*_ value, indicates distinct translation dynamics for these proteins resulting from their increased elongation pausing during aging. These results provide evidence that reduced ribosome concentration in aged brains, likely triggered by aberrant pausing events, might remodel a subset of the proteome independently of transcript levels and regulation (Figure 5C).

## Discussion

Our study offers a comprehensive insight into how distinct mechanisms mediating proteostasis can influence the vertebrate brain proteome during aging. We show significant proteome changes in aging brains, encompassing protein synthesis, solubility, post-translational modifications, and organelle composition. Among these, we show that ribosomal and DNA/RNA binding protein levels decrease independently of mRNA abundance. We propose that translation alterations, including elongation pausing at basic residues, are central drivers of these changes leading to discrepancies between mRNA levels, ribosome occupancy, and protein synthesis.

At least two key implications emerge from our findings. Firstly, basic protein complexes like ribosomes, RNA-binding proteins involved in splicing, RNA and DNA polymerases, and the ones involved in DNA repair experience reduced availability with age. This phenomenon, caused by the presence of basic amino acids in their protein sequences, likely has a direct impact on each step of the gene expression process, and could mechanistically connect proteostasis impairment to other canonical aging hallmarks, including DNA damage, epigenetic alterations (*35*), aberrant splicing (*36*) and reduced RNA polymerase activity (*37*). Of note, individual manipulation of protein biosynthesis, as well as any of these pathways, ameliorates aging phenotypes (*37*–*42*,*43*), further highlighting their central role in the aging process.

A second implication is that aging leads to altered mitochondrial composition, partially driven by a reduced ribosomal content. This remodeling encompasses a decrease of mitochondrial ribosomes, while respiratory chain components remain stable or increase, as in the case of Complex IV. This is consistent with broader observations of aging-induced mitochondrial changes (*44*,*45*). These findings based on bulk tissue measurements were corroborated by more direct analysis of the composition of mitochondria from subcellular fractions and by other age-dependent alterations of mitochondrial proteins, e.g., in detergent insolubility. In addition, we show that a reduced mtDNA content is induced by decreased proteasome activity, showcasing the convergence of aging mechanisms affecting critical cellular structures.

The mechanisms leading to increased translation pausing remain unclear. Future analyses should clarify the mechanics of these events and their relationship with other age-related alterations of ribosomes, including loss of stoichiometry and aggregation (*6*). We speculate that one of the mechanisms contributing to increased translational pausing could reside in the decrease in ATP levels that is typically observed in old tissues (*46*–*49*). This reduction in energy levels might alter the decoding kinetics for specific non-optimal codons, such as those encoding basic amino acids (*50*, *51*), leading to a decreased synthesis rate for these proteins. We also identified distinctive changes in protein ubiquitylation in ribosomal proteins, some of which have been previously associated with ribosome collision induced by different types of translation or proteotoxic stress (*26*, *27*). However, it remains unclear whether these modifications are a cause or consequence of increased pausing. For ribosomes, decoupling in aging manifests as a decrease in protein levels together with a progressive increase in transcript levels. These findings are consistent with several observations. First, an age-dependent increase of transcripts encoding for ribosomal proteins has been observed by single-cell RNAseq in multiple cell types of the murine brain (*52*). Accordingly, increased levels of transcripts encoding for ribosomal proteins were one of the most consistent transcriptional signatures of longevity shared across multiple tissues and mammalian species (*53*). Interestingly, our results suggest that this increase might not result from increased transcription but rather from increased mRNA stability. Decreased abundance of ribosomal proteins with age has been described in multiple organs in mice (*54*), as well as in nematodes (*55*), and the protein half-life of ribosomes is affected by aging in the mouse brain (*56*). These data suggest that similar mechanisms might affect ribosomes in different cell types and organs during mammalian aging. Translation pausing may also represent a converging pathophysiological mechanism shared between aging and neurodegenerative diseases, as ribosome stalling has been linked to perturbation of proteostasis in different types of neurodegenerative diseases (*57*–*60*).

Other mechanisms that have not been investigated in this study can additionally contribute to alteration in proteome composition. For instance, age-dependent impairment of protein degradation by the autophagy-lysosome system can lead to the accumulation of specific proteins (*61*), as has been shown for myelin basic protein (MBP) in microglia (*62*). In addition, stalling of RNA polymerase II has been described to occur with aging, thereby skewing the output of transcription in a gene-length dependent manner (*37*), consistent with a systemic loss of long transcripts observed in multiple aging tissues and species (*63*). A reduction in the abundance of specific transcripts could increase transcriptional noise, lead to an imbalance in the stoichiometry of protein complexes, but also alter the relationship between mRNA and protein levels, especially for long-lived proteins.

Finally, our work might contribute to understanding the relationship between aging and the risk of neurodegenerative diseases. We provide an unprecedented resource (accessible at https://genome.leibniz-fli.de/shiny/orilab/notho-brain-atlas/ credentials username: reviewer password: nothobrain2023) of proteome alterations in the aging vertebrate brain and show that multiple proteins and signaling pathways associated with neurodegeneration in humans become perturbed in different ways during physiological aging in killifish (Supplementary text and Figure S5-S6). Such alterations might underlie convergent mechanisms between aging and mutations that increase the risk of neurodegeneration in old individuals.

## Acknowledgments

The authors gratefully acknowledge support from the FLI Core Facilities Proteomics, Sequencing, and the Fish Facility as well as the Stanford Genomics Facility. A.O. is supported by the German Research Council (Deutsche Forschungsgemeinschaft, DFG) via the Research Training Group ProMoAge (GRK 2155), the Else Kröner Fresenius Stiftung (award number: 2019_A79), the Fritz-Thyssen Foundation (award number: 10.20.1.022MN), the Chan Zuckerberg Initiative Neurodegeneration Challenge Network (award numbers: 2020-221617, 2021-230967 and 2022-250618), and the NCL Stiftung. J.H.L. was supported by the National Institute on Aging of the National Institutes of Health under Award Number T32 AG000266, and research by NIH grants GM05643319 and AG054407 to J.F. A.C. is supported by the German Research Council (Deutsche Forschungsgemeinschaft, DFG, award numbers CE 257/9-1 and CE 257/9-2), by Next Generation EU (PNRR), "Tuscany Health Ecosystem", THE project code ECS 00000017 and the Italian Ministry of University and Research (MIUR) with the program “Joined research for Special Status School”, PRO3. The content is solely the responsibility of the author(s) and does not necessarily represent the official views of the National Institutes of Health. The FLI is a member of the Leibniz Association and is financially supported by the Federal Government of Germany and the State of Thuringia.

## Author contributions

Conceptualization: DDF, AM, JF, AC, AO

Data curation: DDF, AM, JHL, AKS, MT

Investigation: DDF, AM, JHL, EKS, MB, SB

Methodology: DDF, AM, JHL, EKS, MB, SB

Project administration: AO, AC

Data analysis: DDF, AM, JHL, PS, GS

Supervision: ETT, JG, JF, AC, AO

Visualization: DDF, AM, JHL, SB

Writing – original draft: DDF, AM, AO

Writing – review & editing: JHL, EKS, MB, SB, PS, AKS, GS, JF, AC

## Supplementary Materials

### Materials and methods

#### Animal management practices

All experiments were performed in accordance with relevant guidelines and regulations. Fish were bred and kept in FLI’s fish facility according to §11 of the German Animal Welfare Act under license number J-003798. The animal experiment protocols were approved by the local authority in the State of Thuringia (Veterinaer- und Lebensmittelueberwachungsamt; proteasome impairment: reference number 22-2684-04-FLI-19-010). Sacrifice and organ harvesting of non-experimental animals were performed according to §4(3) of the German Animal Welfare Act.

#### *In vivo* proteasome impairment

Adult animals (12–14 wph) were subjected to pharmacological intervention via intraperitoneal injections (IP) during a 4-weeks period of treatment. On each of the sixth day (t = 0, t = 6 d, t = 12d, t = 18d, t = 24d), fish were anesthetized with 200 mg/l buffered MS-222 (PharmaQ) and gently manipulated to deliver IP of Bortezomib at 500 μM or vehicle (1% DMSO in a physiological salt solution) at a dosage of 10 μl/g body weight. Animals from the same hatch were randomly allocated to the experimental groups. Both male and female fish were included in each experimental group. Individual brains from the fish were collected on the last day of treatment and snap-frozen in liquid nitrogen.

#### Proteasome activity assay

CT-L (chymotrypsin-like) proteasome activity was assayed with the hydrolysis of a specific fluorogenic substrate, Suc-LLVY-AMC (UBPBio, Catalog Number G1100). On the day of the experiment, brains were lysed in buffer (50 mM HEPES, pH 7.5 (Sigma Aldrich, H3375); 5 mM EDTA (Carl Roth, 8043.2); 150 mM NaCl (Carl Roth, 3957.1); 1 % (v/v) Triton X-100 (Carl Roth, 3051.3); 2 mM ATP (Sigma Aldrich, A2383) prepared with Milli-Q water) to a final estimated protein concentration of ∼4 mg/mL and homogenized by sonication (Bioruptor Plus) for 10 cycles (30 sec ON/60 sec OFF) at high setting, at 4°C. Lysates corresponding to 10 μg protein were incubated in 50 mM Tris-HCl, pH 7.4, 5 mM MgCl2, 1 mM ATP, 1 mM DTT, 10% glycerol, and 10 μM proteasome substrate for 1 h at 37 °C. Specific proteasome activity was determined as the difference between the total activity of protein extracts and the remaining activity in the presence of 20 μΜ MG132 (Enzo Life Sciences, BML-PI102-0005). Fluorescence was measured by multiple reads for 60 min at 37°C by TECAN Kinetic Analysis (excitation 380 nm, emission 460 nm, read interval 5 min) on a Safire II microplate reader (TECAN).

#### Sample preparation for total proteome and analysis of PTMs

Snap-frozen brains were thawed and transferred into Precellys® lysing kit tubes (Keramik-kit 1.4/2.8 mm, 2 ml (CKM)) containing 150 μl of PBS supplemented with cOmplete™, Mini, EDTA-free Protease Inhibitor (Roche,11836170001) and with PhosSTOP™ Phosphatase Inhibitor (Roche, 4906837001). Based on estimated protein content (5% of fresh tissue weight), three to six brains were pooled to obtain ∼1.5 mg of protein extract as starting material for each biological replicate. Tissues were homogenized twice at 6000 rpm for 30 s using Precellys® 24 Dual (Bertin Instruments, Montigny-le-Bretonneux, France), and the homogenates were transferred to new 2 ml Eppendorf tubes. Proteins were quantified using Pierce™ BCA Protein Assay Kit (Thermo Scientific, 23225), and 1.25 mg was processed for further analysis. Volumes were adjusted using PBS and one-fourth of the volume equivalent of the 4× lysis (8% SDS, 100 mM HEPES, pH8) buffer was added. Samples were sonicated twice in a Bioruptor Plus for 10 cycles with 1 min ON and 30 s OFF with high intensity at 20 °C. The lysates were centrifuged at 18,407 x*g* for 1 min and transferred to new 1.5 ml Eppendorf tubes. Subsequently, samples were reduced using 10 mM DTT (Carl Roth, 6908) for 15 min at 45 °C and alkylated using freshly made 200 mM iodoacetamide (IAA) (Sigma-Aldrich, I1149) for 30 min at room temperature in the dark. An aliquot of each lysate was used for estimating the precise protein quantity using BCA (Thermo Scientific, 23225). Subsequently, proteins were precipitated using cold acetone, as described in (*64*), and resuspended in 500 µl of digestion buffer (3 M urea, 100 mM HEPES pH 8.0). Aliquots corresponding to 20, 200, and 1000 µg protein were taken for proteome, phosphopeptides, and ubiquitylated/acetylated peptides enrichment, respectively, and digested using LysC 1:100 enzyme:proteins ratio for 4 hours (Wako sequencing grade, 125-05061) and trypsin 1:100 enzyme:proteins ratio for 16 hours (Promega sequencing grade, V5111). The digested proteins were then acidified with 10% (v/v) trifluoroacetic acid and desalted using Waters Oasis® HLB µElution Plate 30 µm (2, 10, and 30 mg, depending on the amount of starting material) following manufacturer instructions. The eluates were dried down using a vacuum concentrator and reconstituted in MS buffer A (5% (v/v) acetonitrile, 0.1% (v/v) formic acid). For PTM enrichment, peptides were further processed as described below. For Data Independent Acquisition (DIA) based analysis of total proteome, samples were transferred to MS vials, diluted to a concentration of 1 µg/µL, and spiked with iRT kit peptides (Biognosys, Ki-3002-2) prior to analysis by LC-MS/MS.

#### Sequential enrichment of ubiquitylated and acetylated peptides

Ubiquitylated and acetylated peptides were sequentially enriched starting from ∼1000 µg of dried peptides per replicate. For the enrichment of ubiquitylated peptides, the PTMScan® HS Ubiquitin/SUMO Remnant Motif (K-ε-GG) kit (Cell Signaling Technology, 59322) was used following manufacturer instructions. The K-ε-GG modified enriched fraction was desalted and concentrated as described above, dissolved in MS buffer A, and spiked with iRT kit peptides prior to LC-MS/MS analysis. The flowthrough fractions from the K- ε -GG enrichment were acidified with 10% (v/v) trifluoroacetic acid and desalted using Oasis® HLB µElution Plate 30 µm (30 mg) following manufacturer instructions. Acetylated peptides were enriched as described by (*65*). Briefly, dried peptides were dissolved in 1000 µl of IP buffer (50 mM MOPS pH 7.3, 10 mM KPO_4_ pH 7.5, 50 mM NaCl, 2.5 mM Octyl β-D-glucopyranoside) to reach a peptide concentration of 1 µg/µL, followed by sonication in a Bioruptor Plus (5 cycles with 1 min ON and 30 s OFF with high intensity at 20 °C). Agarose beads coupled to an antibody against acetyl-lysine (ImmuneChem Pharmaceuticals Inc., ICP0388-5MG) were washed three times with washing buffer (20 mM MOPS pH 7.4, 10 mM KPO4 pH 7.5, 50 mM NaCl) before incubation with each peptide sample for 1.5 h on a rotating well at 750 rpm (STARLAB Tube roller Mixer RM Multi-1). Samples were transferred into Clearspin filter microtubes (0.22 µm) (Dominique Dutscher SAS, Brumath, 007857ACL) and centrifuged at 4 °C for 1 min at 2000 x*g*. Beads were washed first with IP buffer (three times), then with washing buffer (three times), and finally with 5 mM ammonium bicarbonate (three times). Thereupon, the enriched peptides were eluted first in basic condition using 50 mM aqueous NH3, then using 0.1% (v/v) trifluoroacetic acid in 10% (v/v) 2-propanol and finally with 0.1% (v/v) trifluoroacetic acid. Elutions were dried down and reconstituted in MS buffer A (5% (v/v) acetonitrile, 0.1% (v/v) formic acid), acidified with 10% (v/v) trifluoroacetic acid, and then desalted with Oasis® HLB µElution Plate 30 µm. Desalted peptides were finally dissolved in MS buffer A, spiked with iRT kit peptides and analyzed by LC-MS/MS.

#### Enrichment of phosphorylated peptides

Lysates (corresponding to ∼200 µg of protein extract) were acetone precipitated, digested into peptides, and desalted, as described in ‘‘Sample preparation for total proteome and analysis of PTMs’’. The last desalting step was performed using 50 μl of 80% ACN and 0.1% TFA buffer solution. Before phosphopeptide enrichment, samples were filled up to 210 µl using 80% ACN and 0.1% TFA buffer solution. Phosphorylated peptides were enriched using Fe(III)-NTA cartridges (Agilent Technologies, G5496-60085) in an automated fashion using the standard protocol from the AssayMAP Bravo Platform (Agilent Technologies). In short, Fe(III)-NTA cartridges were first primed with 100 µl of priming buffer (100% ACN, 0.1% TFA) and equilibrated with 50 μL of buffer solution (80% ACN, 0.1% TFA). After loading the samples into the cartridge, the cartridges were washed with an OASIS elution buffer, while the syringes were washed with a priming buffer (100% ACN, 0.1% TFA). The phosphopeptides were eluted with 25 μL of 1% ammonia directly into 25 μL of 10% FA. Samples were dried down with a speed vacuum centrifuge and stored at −20 °C until LC-MS/MS analysis.

#### Subcellular fraction of killifish brain by LOPIT-DC

All the following steps were performed at 4°C, keeping samples on ice unless stated otherwise. Fresh brains from adult (12 wph) and old (39 wph) killifish were pooled to reach ∼150 mg of wet tissue weight per biological replicate. A mixture of male and female fish was used. Fresh brain tissue was subsequently transferred to a 15 mL Potter homogenizer (Fisher Scientific, 15351321) together with 7.5 mL of lysis buffer (LB) (250 mM sucrose, 10 mM HEPES ph 8.0, 2 mM MgAc, 2 mM EDTA) supplemented with Protease Inhibitor (Roche,11836170001) and homogenized with ∼60 gentle strokes. The brain homogenate was then transferred in a 15mL Falcon tube and treated with Benzonase (Merk, 70664) for 20 min at room temperature. An aliquot of 2.5 mL homogenate was collected for each sample and stored at -80°C to be later processed for differential detergent extraction (see below). The remaining 5 mL were transferred to a 5 mL Eppendorf tube and centrifuged at 500 x*g* for 5 min at 4°C to remove cell debris and unlysed cells. Subsequently, the clarified homogenate was centrifuged at 1000 x*g* for 13 min at 4°C and the resulting pellet was collected as the first subcellular fraction (01). Following one additional centrifugation at 1000 x*g* for 7 minutes, the supernatant was then divided into 4 x 1.5 mL Ultracentrifuge Tubes (Beckman) and processed for differential ultracentrifugation step with an Optima TLX-BenchTop Ultracentrifuge (Beckman, 8043-30-1197), using a TLA55 rotor (Beckman, 366725), using the following ultracentrifugation settings (Table 1):

**Table 1:**
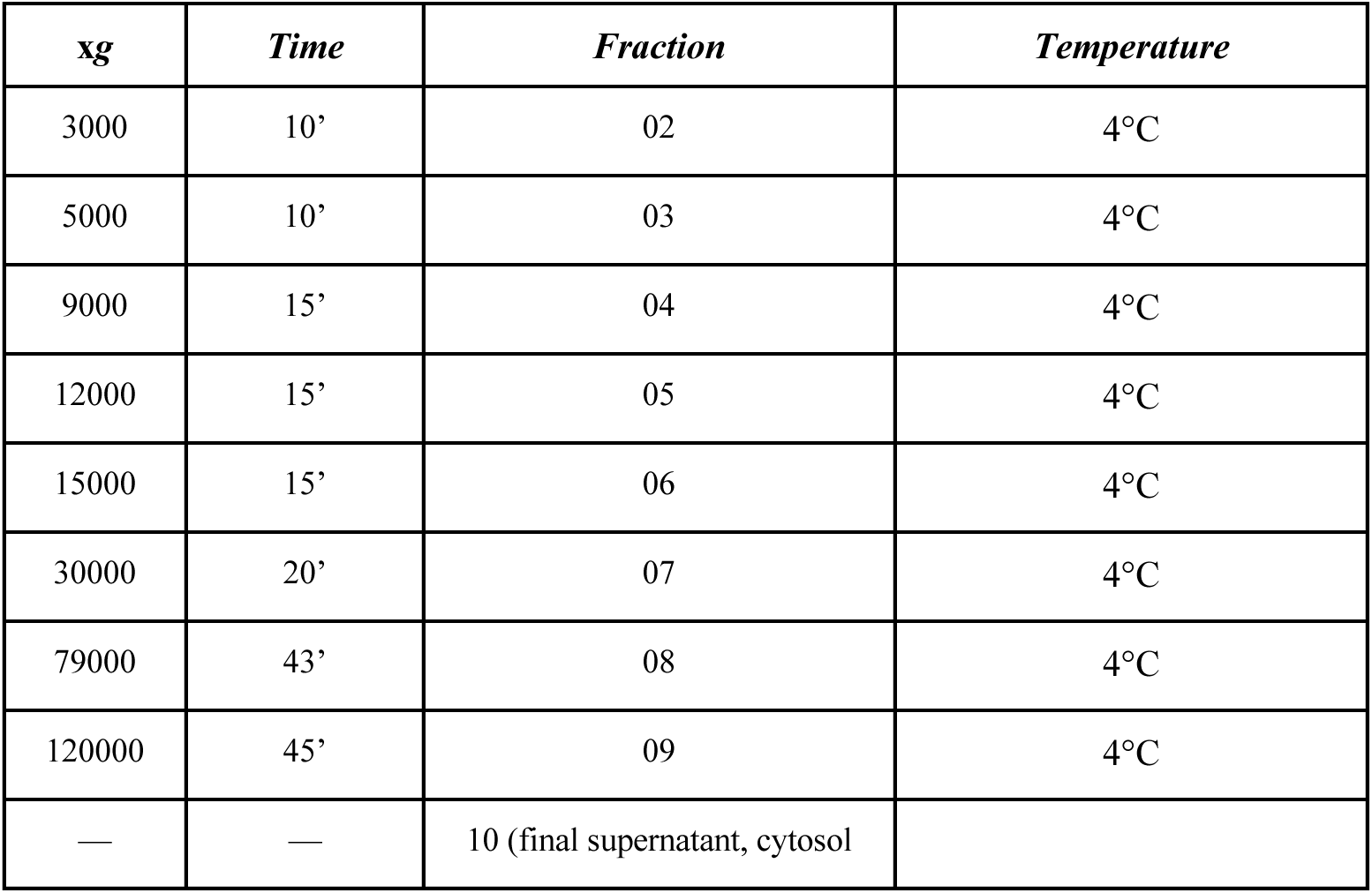
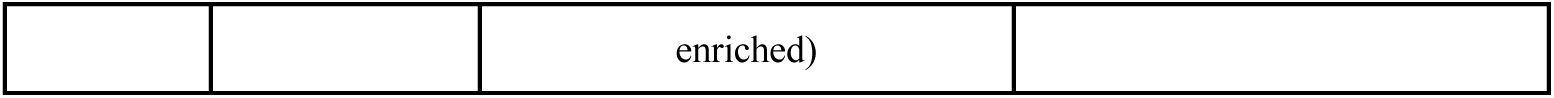
Ultracentrifugation settings for LOPIT-DC protocol.

Pellets from each centrifugation step were resuspended in 50 μL of PBS, and proteins were solubilized by adding 50 μL of 2x lysis buffer (200 mM HEPES pH 8.0, 100 mM DTT, 4% (w/v) SDS). For fraction 10 (cytosol enriched), 300μL was taken and supplemented with 300 μL of 2x lysis buffer. All the samples were then sonicated using a Bioruptor Plus (Diagenode) for 5 cycles with 60 sec ON and 30 sec OFF with max intensity, boiled for 10 min at 95°C, and a second sonication cycle was performed. The solubilized proteins were reduced with 200mM DTT for 15 min at 45°C and alkylated using freshly made 200mM IAA for 30 min at room temperature in the dark. Subsequently, proteins were precipitated using cold acetone, dissolved in 1 M guanidine HCl in 100 mM HEPES pH8.0, and digested using LysC and trypsin, as described in (*64*). The digested proteins were then acidified with 10 % (v/v) trifluoroacetic acid and desalted using Oasis® HLB μElution Plate 30 μm following manufacturer instructions. The eluates were dried down using a vacuum concentrator and reconstituted in 5 % (v/v) acetonitrile, 0.1 % (v/v) formic acid. Samples were transferred directly to MS vials, diluted to a concentration of ∼1 μg/μL, and spiked with iRT kit peptides prior to analysis by LC-MS/MS.

#### Differential detergent extraction

All the following steps were performed at 4°C, keeping samples on ice unless stated otherwise. For each replicate, 2.5 mL of brain homogenate was thawed on ice. After thawing, the homogenate was centrifuged at 500 x*g* for 5 min at 4°C to remove debris. The supernatant was collected, and 64 μL of 20% (v/v) IGEPAL Nonidet P-40 (Sigma) was added to reach an initial concentration of 0.5% (v/v). The homogenate was then divided into 4x 1.5mL ultracentrifuge tubes and sonicated in a Bioruptor Plus for 10 cycles with 30 min ON and 30 s OFF with max intensity at 24 °C. The homogenates were then loaded into a TLA55 rotor and ultracentrifuged with an Optima TLX-BenchTop Ultracentrifuge at 100,0000 x*g* for 5 min at 24°C. After ultracentrifugation, the supernatants were collected and stored as “soluble” (S) fraction. The remaining pellets were resuspended in 1mL of buffer A (10 mM HEPES pH 8.0, 2 mM MgAc, 2 mM EDTA, 0.5% NP-40), samples were mixed by vortexing, and sonicated in a Bioruptor Plus for 10 cycles with 30 s ON and 30 s OFF with max intensity at 24 °C. Samples were then ultracentrifuged again at 100,0000 x*g* for 5 min at 24°C. The supernatants (“F1”) were collected and the remaining pellets were resuspended in 1mL of buffer B (10 mM HEPES pH 8.0, 2 mM MgAc, 2 mM EDTA, 0.5% NP-40, 0.25% SDS, 0.5% deoxycholic acid), mixed, sonicated, and centrifuged as above. The supernatants (“F2”) were collected and the remaining pellets were resuspended in 1mL of buffer C (10 mM HEPES pH 8.0, 2 mM MgAc, 2 mM EDTA, 0.5% NP-40, 2% SDS, 0.5% deoxycholic acid), mixed, sonicated, and centrifuged as above. The supernatants (“F3”) and the remaining pellets were collected. All the collected samples were stored at -80°C until further analysis.

#### Data independent acquisition for proteome quantification

Peptides were separated in trap/elute mode using the nanoAcquity MClass Ultra-High Performance Liquid Chromatography system (Waters, Waters Corporation, Milford, MA, USA) equipped with trapping (nanoAcquity Symmetry C18, 5 μm, 180 μm × 20 mm) and an analytical column (nanoAcquity BEH C18, 1.7 μm, 75 μm × 250 mm). Solvent A was water and 0.1% formic acid, and solvent B was acetonitrile and 0.1% formic acid. 1 μl of the samples (∼1 μg on column) were loaded with a constant flow of solvent A at 5 μl/min onto the trapping column. Trapping time was 6 min. Peptides were eluted via the analytical column with a constant flow of 0.3 μl/min. During the elution, the percentage of solvent B increased nonlinearly from 0–40% in 120 min. The total run time was 145 min, including equilibration and conditioning. The LC was coupled to an Orbitrap Exploris 480 (Thermo Fisher Scientific, Bremen, Germany) using the Proxeon nanospray source. The peptides were introduced into the mass spectrometer via a Pico-Tip Emitter 360-μm outer diameter × 20-μm inner diameter, 10-μm tip (New Objective) heated at 300 °C, and a spray voltage of 2.2 kV was applied. The capillary temperature was set at 300°C. The radio frequency ion funnel was set to 30%. For DIA data acquisition, full scan mass spectrometry (MS) spectra with a mass range 350–1650 m/z were acquired in profile mode in the Orbitrap with the resolution of 120,000 FWHM. The default charge state was set to 3+. The filling time was set at a maximum of 60 ms with a limitation of 3 × 10^6^ ions. DIA scans were acquired with 40 mass window segments of differing widths across the MS1 mass range. Higher collisional dissociation fragmentation (stepped normalized collision energy; 25, 27.5, and 30%) was applied, and MS/MS spectra were acquired with a resolution of 30,000 FWHM with a fixed first mass of 200 m/z after accumulation of 3 × 10^6^ ions or after filling time of 35 ms (whichever occurred first). Data were acquired in profile mode. For data acquisition and processing of the raw data, Xcalibur 4.3 (Thermo) and Tune version 2.0 were used.

#### Data processing for MS-DIA samples

Spectral libraries were created by searching the DIA or/and DDA runs using Spectronaut Pulsar (14.9.2 and 15.3.2, Biognosys, Zurich, Switzerland). The data were searched against species-specific protein databases (Nfu_20150522, annotation nfurzeri_genebuild_v1.150922) with a list of common contaminants appended. The data were searched with the following modifications: carbamidomethyl (C) as fixed modification, and oxidation (M), acetyl (protein N-term), lysine di-glycine (K-ε-GG), phosphorylated tyrosine (T) and serine (S) and acetyl-lysine (K-Ac) as variable modifications for the respective PTMs enrichments. A maximum of 3 missed cleavages were allowed for K-Ac and K-ε-GG modifications, 2 missed cleavages were allowed for phospho enrichment. The library search was set to 1 % false discovery rate (FDR) at both protein and peptide levels. DIA data were then uploaded and searched against this spectral library using Spectronaut Professional (v14.9.2 and 15.3.2) and default settings. Relative quantification was performed in Spectronaut for each pairwise comparison using the replicate samples from each condition using default settings, except the one displayed in Table 2:

**Table 2:** Setting list used for MS data analysis on Spectronaut Software.

| *Dataset* | *Software version* | *Test* | *Data Filtering* | *Imputation* | *Normalization* |
| --- | --- | --- | --- | --- | --- |
| Aging proteome | 15.3.2 | Unpaired t-test | Q-value | Global Imputing | True, Automatic |
| LOPIT-DC | 14.9.2 | NA | Q-value percentile 0.2 | Run Wise Imputing | True, Global |
| Detergent insolubility | 15.4.2 | NA | Q-value percentile 0.2 | Run Wise Imputing | False |
| Proteasome Inhibition | 14.9.2 | Unpaired t-test | Q-value | Global Imputing | True, Automatic |
| PTMs - Ubiquitin | 15.4.2 | – | Q-value percentile 0.2 | Global Imputing | True, Automatic |
| PTMs - Phosphorylation | 15.4.2 | – | Q-value percentile 0.2 | Global Imputing | True, Automatic |
| PTMs - Acetylation | 15.4.2 | – | Q-value percentile 0.2 | Global Imputation | True, Automatic |

Candidates and report tables were exported from Spectronaut and used for downstream analysis.

#### Immunoblot

Killifish brains and cells treated for 24 hours with anisomycin (Cell Signaling Technology, 2222) were lysed following as described in “Sample preparation for total proteome and analysis of PTMs”. Protein concentration was estimated by Qubit assay (Invitrogen, Q33211), and 30 µg of proteins were used. 4× loading buffer (1.5 M Tris pH 6.8, 20% (w/v) SDS, 85% (v/v) glycerin, 5% (v/v) β-mercaptoethanol) was added to each sample and then incubated at 95 °C for 5 minutes. Proteins were separated on 4–20% Mini-Protean® TGX™ Gels (BioRad, 4561096) by sodium dodecyl sulfate-polyacrylamide gel electrophoresis (SDS-PAGE) using a Mini-Protean® Tetra Cell system (BioRad, Neuberg, Germany, 1658005EDU). Proteins were transferred to a nitrocellulose membrane (Carl Roth, 200H.1) using a Trans-Blot® Turbo™ Transfer Starter System (BioRad, 1704150). Membranes were stained with Ponceau S (Sigma, P7170-1L) for 5 min on a shaker (Heidolph Duomax 1030), washed with Milli-Q water, imaged on a Molecular Imager ChemiDocTM XRS + Imaging system (BioRad) and destained by 2 washes with PBS and 2 washes in TBST (Tris-buffered saline (TBS, 25 mM Tris, 75 mM NaCl), with 0.5% (v/v) Tween-20) for 5 min. After incubation for 5 min in EveryBlot blocking buffer (Biorad, 12010020), membranes were incubated overnight with primary antibodies against RPS3 (Bethyl Laboratories, A303-840A-T) or α-tubulin (Sigma, T9026) diluted (1:1000) in enzyme dilution buffer (0.2% (w/v) BSA, 0.1% (v/v) Tween20 in PBS) at 4 °C on a tube roller (BioCote® Stuart® SRT6). Membranes were washed 3 times with TBST for 10 min at room temperature and incubated with horseradish peroxidase coupled secondary antibodies (Dako, P0448/P0447) at room temperature for 1 h (1:2000 in 0.3% (w/v) BSA in TBST). After 3 more washes for 10 min in TBST, chemiluminescent signals were detected using ECL (enhanced chemiluminescence) Pierce detection kit (Thermo Fisher Scientific, Waltham, MA, USA, #32109). Signals were acquired on the Molecular Imager ChemiDocTM XRS + Imaging system and analyzed using the Image Lab 6.1 software (Biorad). Membranes were stripped using stripping buffer (1% (w/v) SDS, 0.2 M glycine, pH 2.5), washed 3 times with TBST, blocked, and incubated with the second primary antibody, if necessary.

#### RNA isolation for RNA-Seq analysis

Individual brains from the fish were collected and snap-frozen in liquid nitrogen. The protein amount was estimated based on fresh tissue weight (assuming 5% of protein w/w), and ice-cold 1x PBS with protease/ phosphatase inhibitors (Roche,11836170001, 4906837001) was added accordingly to a final concentration of 2 μg/μL. Samples were then vortexed (5 times) before sonication (Bioruptor Plus) for 10 cycles (60 sec ON/30 sec OFF) at the high setting, at 4 °C. The samples were then centrifuged at 3000 x*g* for 5 min at 4 °C, and the supernatant was transferred to 2 mL Eppendorf tubes. 1.5 mL of ice-cold Qiazol (Qiagen, 79306) reagent was added to 150 μL of homogenate, vortexed five times, and snap-frozen in liquid nitrogen. On the day of the experiment, samples were thawed on ice, vortexed five times, and incubated at room temperature for 5 min before adding 300 μL of chloroform. Samples were mixed vigorously, incubated for 3 min at room temperature, and centrifuged at 12000 x*g* for 20 min at 4 °C. The upper aqueous phase (600 μL) was carefully transferred into a fresh tube, and the remaining volume (phenol/chloroform phase) was kept on ice for DNA isolation. The aqueous phase was mixed with 1.1 volume of isopropyl alcohol, 0.16 volumes of sodium acetate (2 M; pH 4.0), and 1 μL of GlycoBlue (Invitrogen, AM9515) to precipitate RNA. After 10 min incubation at room temperature, samples were centrifuged at 12000 x*g* for 30 min at 4 °C. The supernatant was completely removed, and RNA pellets were washed by adding 80% (v/v) ethanol and centrifuging at 7500 x*g* for 5 min at 4 °C. The washing steps were performed twice. The resulting pellets were air-dried for no more than 5 min and dissolved in 10 μL nuclease-free water. To ensure full dissolution of RNA in water, samples were then incubated at 65 °C for 5 min, before storage at -80 °C.

#### RNA-Seq library preparation

Sequencing of RNA samples was done using Illumina’s next-generation sequencing methodology (*66*). In detail, quality check and quantification of total RNA was done using the Agilent Bioanalyzer 2100 in combination with the RNA 6000 pico kit (Agilent Technologies, 5067-1513). Total RNA library preparation was done by introducing 500 ng total RNA into Illumina’s NEBNext Ultra II directional mRNA (UMI) kit (NEB, E7760S), following the manufacturer’s instructions. The quality and quantity of all libraries were checked using Agilent’s Bioanalyzer 2100 and DNA 7500 kit (Agilent Technologies, 5067-1506).

#### RNA-Seq sequencing

All libraries were sequenced on a NovaSeq6000 SP 300 cycles v1.5; paired-end 151 bp (one pair for each of the projects). Total RNA libraries were pooled and sequenced in three lanes. Small RNA libraries were pooled and sequenced in one lane. Sequence information was extracted in FastQ format using Illumina’s bcl2FastQ v2.20.0.422, against the *Nothobranchius furzeri* reference genome (Nfu_20150522, annotation nfurzeri_genebuild_v1.150922). Alignment to the reference genome was performed using STAR (*67*) with the following parameters: --outSAMmultNmax 1 -- outFilterMultimapNmax 1 -- outFilterMismatchNoverLmax 0.04 --sjdbOverhang 99 --alignIntronMax 1000000 -- outSJfilterReads Unique. The deduplication step was performed using the umi_tool v1.1.1 (*68*), using the following parameters: extract --bcpattern= NNNNNNNNNNN’, ‘dedup --chimeric-pairs discard --unpaired-reads discard -- paired.

#### RNA-Seq quantification and differential expression

RNA-Seq data were then processed as follows: quantification was performed using featurecounts v2.0.3 (*69*) with the following parameters -s 2 -p -B --countReadPairs. Differential expression analysis was performed using the DESeq2 package (v1.34.0) (*70*). Raw count data were normalized using the transcript per million strategy.

#### Ribo-Seq library preparation

Ribosome profiling libraries were prepared following previously published protocol with modifications (*24*). 10∼15 brain samples from fish were combined and lysed frozen using Cryo-Mill (Retsch, MM301) in the presence of 1ml of lysis buffer (20 mM Tris-HCl pH 7.5, 140 mM KCl, 5 mM MgCl2, 1 mM DTT, 100 µg/ml Cycloheximide, 1% Triton X-100, and 1 X Protease Inhibitor). Lysed powder was quickly thawed in a water bath at room temperature and spun at 21,000 g for 15 minutes at 4 °C to clear lysate. RNAse I (Invitrogen, AM2294) was added to 0.4U/μg of RNA and incubated at 25 °C for 45 minutes. Digestion was stopped by adding 0.4U/μg of SUPERaseIn RNAse Inhibitor (Invitrogen, AM2696). RNAse-treated lysate was layered on 900 μl sucrose cushion buffer (20 mM Tris-HCl pH 7.5, 140 mM KCl, 5 mM MgCl2, 1 mM DTT, 100 µg/ml Cycloheximide, 0.02U/μl SuperaseIn, 1M Sucrose), and spun at 100,000 rpm for 1 hour at 4 °C in TLA100.3 rotor. Resulting ribosome pellet was resuspended in 250 μl of lysis buffer with SuperaseIn and RNA was extracted using TRIzol reagent (Invitrogen, 15596026) following manufacturer’s protocol. 27-34bp fragments were isolated from denaturing gel, ligated to adapter (NEB, S1315S), and ribosomal RNA was removed using RiboCop (Lexogen, 144.24) mixed with custom depletion DNA oligos (Table 4). Remaining fragments were reverse transcribed, circularized, and PCR amplified following the steps described previously (*71*). Barcoded samples were pooled and sequenced using Hiseq 4000 (Illumina).

#### Imaging

##### Cryo-sections preparation and free-floating immunofluorescence

To prepare brain cryo-sections for free-floating immunofluorescence from 5 wph and 39 wph old killifish, brains were dissected and fixed ON in a solution of 4% paraformaldehyde PFA in PBS at 4°C. The samples were then equilibrated in a 30% sucrose solution ON at 4° and subsequently embedded in cryo-protectant (Tissue -Tek O.C.T. Compound; Sakura Finetek, USA). Tissue slices of 50mm thickness were cut at a cryostat (Leica) and stored on glass slides (Thermo Fisher Scientific, USA).

Free-floating immunofluorescence experiments were performed by adapting previous protocols for classical on-slide immunofluorescence (*72*). Briefly, the sections were washed in PBS to remove the cryo-embedding medium and detached from the glass slide. The sections were then placed in 24-wells and performed two additional washes in PBS for 5 min each. Afterward, an acid antigen retrieval step (10 mM Tri-sodium citrate dihydrate, 0.05% tween, at pH 6) was performed by bringing the solution to boiling point in a microwave and adding 50ml of it in each well, leaving the solution for 5 minutes. This step was repeated two times.. 500 ml of blocking solution (5% BSA, 0.3% Triton-X in PBS) was then applied for 2 h. Primary antibodies (Phospho-Tau AT100, NeuN or Lamp1 Table 3) at the proper dilution were added in a solution of 1% BSA, 0.1% triton in PBS, and left overnight at 4°C in slow agitation on a rocker. Next day, the proper secondary antibodies (Table 3) at a 1:500 dilution were used in the same solution. After 2h of incubation, slices were washed three times with PBS, counter-stained with a solution 1:10000 of Hoechst 33342 (Invitrogen, USA) for two minutes and manually mounted under a stereomicroscope on Superfrost Plus glass slides (Thermo Fisher Scientific, USA). Finally, Fluoroshield mounting medium (Sigma, USA) was used and slices were covered with a coverglass (Thermo Fisher Scientific, USA).

**Table 3:** List of antibodies utilized in this work.

| Primary Antibody | Producer | Catalog Number | Type | Working dilution |
| --- | --- | --- | --- | --- |
| Lamp1 | Abcam | Ab24170 | Polyclonal Rabbit | 1:500 |
| NeuN | Abcam | Ab177487 | Monoclonal Rabbit | 1:500 |
| Phospho-Tau AT100 | Thermo Fisher Scientific | MN1060 | Monoclonal Mouse | 1:400 |
| Secondary Antibody |  |  |  |  |
| AlexaFluor 488 anti-Rabbit | Invitrogen | A11001 | Goat IgG | 1:500 |
| AlexaFluor 568 anti-Rabbit | Invitrogen | A11011 | Goat IgG | 1:500 |
| AlexaFluor 488 anti-Mouse | Invitrogen | A11004 | Goat IgG | 1:500 |

**Table 4:**
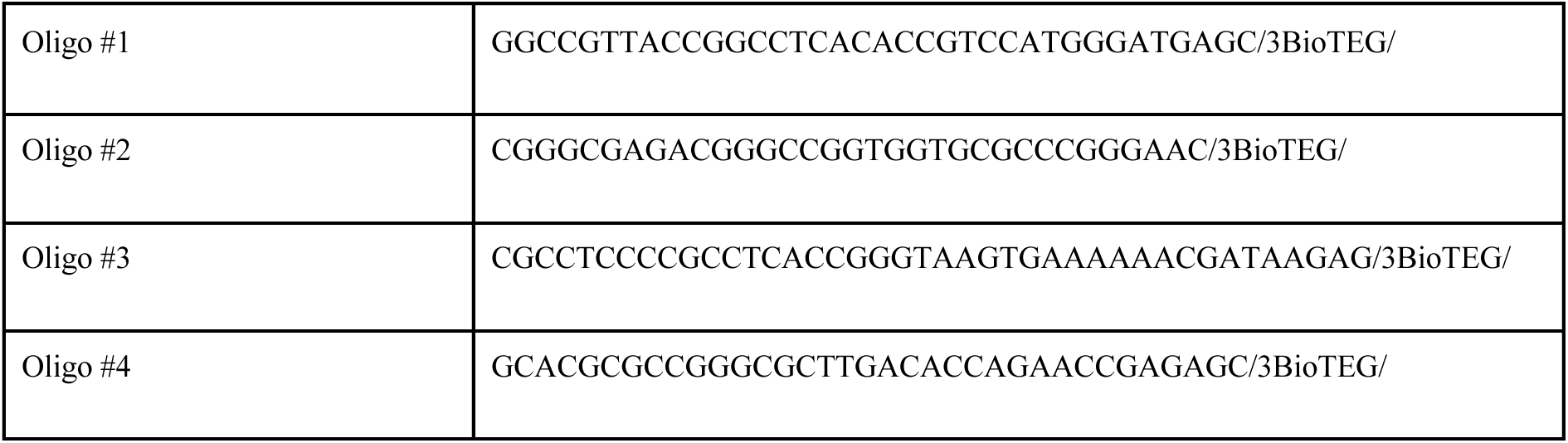
List of DNA oligonucleotides used for ribosomal RNA depletion.

##### Image acquisition

Imaging of lysosomal staining was performed with a Zeiss scanning confocal microscope (LSM900, Zeiss, Germany) equipped with an Airyscan module. Nine consecutive z planes with a step of 300nm were acquired with a 63x oil immersion objective (Plan-Apochromat 63x/1.4 Oil DIC M27, Zeiss, Germany) at a resolution of 2186x2186 pixels with the use of Airyscan. Images were then deconvoluted in the Zeiss Zen blue 3.7 suite using the Fast Iterative algorithm and exported as tiff for further analysis in Imaris (Bitplane, UK).

Samples processed for Tau stainings were imaged with an Axio Imager Z.2 (Zeiss, Germany) equipped with an Apotome slide using a 63x oil immersion objective (Plan-Apochromat 63x/1.4 Oil DIC M27, Zeiss, Germany). Z-stacks were realized by acquiring five consecutive z-planes at an interval of 1 micron. Images were then processed in imageJ (Fiji).

##### Lysosomes morphological analysis

To analyze the change in morphology of lysosomes in aging, we analyzed nine 5 wph samples and twelve 39 wph samples. To study morphological changes in case of proteostasis alteration, samples from six bortezomib-treated animals and six controls (DMSO treated) were analyzed. Tiff images were loaded in Imaris (Bitplane, UK) to recreate a 3D rendering of the samples. A version of the ‘Surfaces’ algorithm was created, optimizing the settings to realize an optimal mask of single lysosomes. Statistics obtained (Area, Volume, Mean intensity, and Sphericity) were extracted, and mean values for each animal were calculated. Data significance was tested using a two-tails T-test.

##### Mean fluorescence intensity analysis

To analyze differences in the amount of Tau phosphorylation between young (5 wph) and old (39 wph) *Nothobranchius furzeri* brain samples, we performed mean fluorescence intensity (MFI) analysis in the free license software ImageJ (Fiji). Since Tau is a neuronal protein, and the number of neurons between young and old animals varies, we normalized the MFI of Tau staining over the MFI of NeuN, a neuronal-specific marker, in order to render the Tau MFI proportional to the number of neurons. Images were opened in ImageJ (Fiji), and median filtering (1px radius) was applied. The average intensity projection was realized, and MFI for the green channel (Tau) and red channel (NeuN) was measured and reported in an Excel table. Tau MFI for each animal was divided by the corresponding NeuN MFI, and the significance of the results was tested by a two-tails T-test.

### Data analysis

#### Protein subcellular localization by LOPIT-DC

For each age group and replicate, protein distribution profiles were calculated by dividing the scaled protein quantity in each fraction by the total sum of protein quantity across all fractions. Protein markers for the different compartments were taken from the Bioconductor package pRoloc (*73*), by mapping *Nothobranchius furzeri* entries onto *Homo sapiens* entries via orthologues mapping. To classify each of the proteins into a stable compartment, a support-vector-machine classifier with a radial kernel (*74*) was used. Hyper-parameters *C* and *gamma* were selected via a grid-search approach using a 5-fold cross-validation iterated 100 times. The best *C* and *gamma* parameters were selected to classify the “unknown” proteome. Only classified proteins with an SVM-score > 0.7 were considered stable classification. To detect age-related changes in subcellular fractionation, a two-step approach was implemented. For each normalized protein profile, a principal component analysis was used to summarize the variance from the 10 fractions in each replicate and age group. After summarization, the first two principal component scores were used to perform a Hotelling T^2^ test to detect changes in the multivariate protein profile mean. To estimate effect sizes, the median Euclidean distance between age groups was calculated for each protein profile (see Figure S3F).

#### Differential detergent extraction

A batch correction was applied to remove the effects of different batches of LC-MS/MS analysis using the limma::removeBatchEffect function from the limma package (*75*). Then, for each protein group, a detergent insolubility profile was generated by dividing the protein quantities from fractions F1:F3 by the quantity in the soluble (S) fraction, and log2 transformed. To detect significant changes in detergent insolubility profiles between age groups, a MANOVA test was applied to the detergent insolubility profiles using the standard function in the R programming language, and P-values were corrected for multiple testing using the FDR strategy. To estimate effect sizes, a detergent-insolubility-score (DIS) was calculated by summing the log2 transformed protein quantities in fractions F1:F3 relative to the S “soluble” fraction. For each age group and protein group, the median DIS between replicates was used to estimate the magnitude of changes in detergent insolubility: ΔDIS = DIS_39wph_ - DIS_12wph_. High values of ΔDIS indicate proteins that become more detergent resistant in the old (39 wph) samples (see Figure S2F).

#### Modified peptide abundance correction

For each enrichment, PTMs report tables were exported from Spectronaut. To correct the quantities of modified peptides for underlying changes in protein abundance across the age groups compared, correction factors were calculated using the aging proteome data. For each condition and protein group, the median protein quantity was calculated and then divided by the median protein quantity in the young (5 wph) age group. Each modified peptide was matched by protein identifier to the correction factor table. If a modified peptide was mapped to 2 or more proteins, the correction factor was calculated using the sum of the quantity of these proteins. Further, the correction was carried out by dividing peptide quantities by the mapped correction factors, and log2 transformed (see Figure S4B). Differences in peptide quantities were statistically determined using the t-test moderated by the empirical Bayes method as implemented in the R package limma (*75*).

#### Kinase activity prediction from phosphoproteome data

Kinase activity prediction was calculated using the Kinase library (https://kinase-library.phosphosite.org/ea?a=de, (*76*) using the differential expression-based analysis and default parameter.

#### GO enrichment analysis

Gene Set Enrichment Analysis (GSEA) was performed using the R package clusterProfiler (*77*), using the function gseGO. Briefly, *Nothobranchius furzeri* protein entries were mapped to the human gene name orthologues and given in input to the function to perform the enrichment. For GO term overrepresentation analysis (ORA), the topGO R package was used.

#### Identification of conserved PTMs sites

For the *Nothobranchius furzeri* proteins involved in neurodegenerative diseases (Figure S5I), a local alignment was performed with protein BLAST(v2.12.0+) (*78*) with default parameters against the RefSeq human proteome (Taxon ID:9606). The top 10 hits from the BLAST search were retrieved, and each modified residue was mapped into the local alignment to identify the corresponding position in the human proteins. Each modified peptide was then considered conserved if at least one of the top 10 hits from the BLAST alignment had a corresponding residue in the modified amino acid position.

#### Calculation of protein-transcript decoupling and multiple linear regression

For aging brain proteome data and proteasome impairment samples, protein-transcript decoupling values were calculated as the difference in log2 fold changes between proteome and transcriptome. A null distribution was fitted on the decoupling values using the R package fdrtool (*79*). Q-value < 0.1 was used as a threshold to reject the null hypothesis. The decoupling values from each protein-transcript pair were used as response variables in a multiple linear regression model. Predictors for the model were retrieved as follows: protein quantities were calculated as the median log2 protein quantity across all replicates from the proteomics DIA data. Protein quantities are estimated using the median peptide abundance as calculated by the Spectronaut software. mRNA abundance values were defined as the median log2(TPM) across all samples from the RNA-Seq aging dataset. Biophysical parameters were calculated for each protein with the R package Peptides. Protein half-life values were taken from mouse cortex data from (*16*). The percentage of gene GC content was obtained from ENSEMBL Biomart (v108) (*80*), mapping ENSEMBL annotation against the *Nothobranchius furzeri* reference genome (Nfu_20150522, annotation nfurzeri_genebuild_v1.150922) using bedtools (*81*). Multiple linear regression models were then performed using the ‘lm’ base R function by keeping only complete and unique observations from the matrix generated. Features were scaled for each dataset, and a multiple linear regression model without intercept was fitted to the data.

#### Data integration

Log2 fold changes (for PTMs), ΔDIS (for detergent insolubility), or protein-transcript decoupling score values were used as input for a GSEA analysis based on GO cellular component terms using the gseGO function from the clusterProfile (*77*) R package with the following parameters minSize = 5 and maxSize = 400. For each GSEA, the normalized enrichment scores (NES) were taken and arranged in a matrix with different GO terms as rows and different datasets as columns. To visualize the relationship between the dataset, a principal component analysis was performed on the matrix. Missing GO terms in a given dataset were imputed as 0 values. The sum of the scores on the first two principal components was used to extract the most strongly affected GO terms from the combined integration of all the datasets.

#### Mitochondrial proteome composition

To calculate age-related changes in mitochondrial proteome composition (Figure 2H), raw DIA files coming from fraction 02 of the LOPIT-DC experiment were re-analyzed in Spectronaut (v16.2), using the same parameters as the other LOPIT-DC experiment. Fraction 02 represents the fraction where mitochondrial proteins are sedimenting in the LOPIT-DC experiment and, therefore, strongly enriched for mitochondrial proteins (Figure S3C-D). From the protein quantity matrix, mitochondrial proteins (according to Mitocarta3.0 annotation (*82*)) were extracted, and their quantities log2 transformed and normalized by median centering. To detect changes in composition, a linear model on the log2 mitochondrial-centered values was implemented between the two age groups with the R package limma (*75*).

#### Ribo-seq data processing and analysis

Data processing and analysis was based on previously published protocol (*24*). Adapter sequences were removed from demultiplexed sequencing reads using Cutadapt v.1.4.2 (*83*), followed by removal of the 5’ nucleotide using FASTX-Trimmer. Reads mapping to ribosomal RNAs were removed using Bowtie v.1.3.1 (*84*). Remaining reads were aligned to reference libraries that consisted of coding sequences containing 21 nucleotides flanking upstream of the start codon and downstream of the stop codon. To maximize unique mapping, a reference library was constructed using the longest transcripts for every 22757 genes. Bowtie alignment was performed using the following parameters: -y -a -m 1 -v 2 -norc - best -strata. A-site offset was estimated using riboWaltz (*85*), and fragment lengths that do not exhibit 3-nucleotide periodicity were removed. Pause scores at each position were calculated by dividing the number of reads at each position by the average number of reads within the internal part of the transcript, excluding the first and last 20 codons. Positions with increased pausing during aging were identified following the previously published method (*24*). Briefly, for 6749 transcripts with sufficient coverage (>0.5 reads/codon and >64 reads/transcript) in all age groups, we used a two-tailed Fisher’s exact test to compare each position (codon) between age groups to identify positions with statistically significant changes (Benjamini-Hochberg adjusted P-value < 0.05). These positions were further filtered to include positions with odds ratio greater than 1, pause score of the older sample greater than the pause score of younger sample, reads in the oldest sample greater than the average number of reads across the transcript, and a position in the internal part of the transcript to only select sites with high-confidence age-dependent changes in pausing. To visualize amino acids enriched in age-dependent pausing sites, we used the weighted Kullback Leibler method (*86*) using the frequency of each amino acid in coding sequences as background. For metagene analysis around age-dependent pausing sites, reads were first aligned to these sites and normalized by dividing reads at each codon by the average reads per codon within the analysis window to control for differences in expression and coverage. Mean and bootstrapped 95% confidence intervals of these normalized values were plotted. Only positions with sufficient coverage (reads/codon>0.5) in the analysis window were included. To identify sites with disome formation, we first identified sites with strong pausing in the old sample (pause score >6). Then, we calculated the average ribosome density of two regions for young and old samples; 1) analysis window (40 codons up/downstream from strong pause site) and 2) between 8 and 12 codons upstream from strong pause site (approximate position of trailing ribosome). Sites with higher ribosome density in 2) were identified as disome sites, and disomes sites unique to old samples were plotted. For comparisons to proteomics data sets, we included all sites with statistically significant changes (Benjamini-Hochberg adjusted P-value < 0.05) and used log2 of pause score ratio (Old/Young). For translation efficiency analysis, RNA-seq data was re-aligned to the same reference library used for Ribo-seq to compare transcript abundance. Changes in translation efficiency were calculated using DESeq2 (*70*), using the following design ∼assay + condition + assay:condition, where assay indicates the different counts from RNA-Seq and Ribo-Seq respectively, and condition indicated the different age groups.

#### Estimates of mRNA half-life variations

Exonic coordinates of protein-coding genes were extracted from the annotation nfurzeri_genebuild_v1.150922. Exonic and intronic read counts were obtained following the procedure suggested by (*31*). To this end, exonic coordinates were flanked on both sides by 10 nt and were grouped by gene. Intronic coordinates were obtained by subtracting the exonic coordinates from the gene-wise coordinates. For each gene, exonic and intronic read counts were obtained using the htseq-count function from HTSeq v2.0.2 (*87*) with the parameter -m set to intersection-strict to consider only reads that strictly fall within an exon or an intron. Additionally, in each sample, genes with less than 10 reads on both exons and introns were ignored (read counts set as missing values) in order to be robust against noisy estimates based on low read counts. Lastly, the log-transformed exonic-to-intronic read count ratio r was computed for each gene and sample as:

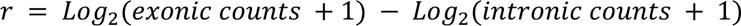

Gene-specific biases such as exonic and intronic lengths and GC content can affect exonic and intronic read counts. These biases cancel out when ratios between samples are considered, as they are typically multiplicative (*31*). The ratio between mRNA half-life in sample s_1 and sample s_2 is then estimated as:

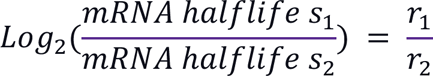

#### Estimates of protein synthesis rate

To estimate *k*_*i*_, 5’-UTRs sequences were retrieved from the *Nothobranchius furzeri* reference genome (Nfu_20150522, annotation nfurzeri_genebuild_v1.150922). The masked FASTA genome sequences were parsed using bedtools (*81*). The translation starting codon “ATG” was identified from the ‘CDS’ features from the GFF file. The region around the starting codon was extracted with +6 nucleotide upstream and +4 nucleotide downstream to match the pattern “NNNNNNATGNN”. Only valid sequences (without ambiguous nucleotides) with an ATG starting codon in the correct position were retained. 91% of the transcript annotated in the GFF file had a valid translation initiation region as described above. The *k*_*i*_ was then estimated using the dinucleotide position weight matrix from (*34*). In case a single transcript had multiple starting sites, the *k*_*i*_ values were summarized by taking the median value. This led to the estimate of *k*_*i*_ for 59129 transcripts. Estimated protein synthesis rates were calculated as in (*32*, *33*). More in detail, the authors described the estimated synthesis rate as:

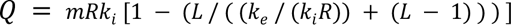

where *Q* refers to the estimated synthesis rate, *m* refers to individual mRNA expression level obtained from the median across sample log2(TPM) from RNA-Seq data and normalized between 0 and 1, *R* represents the total amount of available ribosomes, *k*_*i*_ indicates an mRNA-specific translation initiation rate as computed above and normalized between 0 and 1, *L* is the number of codons occupied by one ribosome, set to 10 (based on the average length of a ribosome footprint), and *k*_*e*_ is the termination rates arbitrarily set to 1. Estimated synthesis rates were then computed for different values of *R* ranging from 1.3 to 0.

#### Supplementary text

Aging can influence different aspects of protein homeostasis. To obtain an unbiased characterization of the effect of aging on the brain proteome we employed a multi-layered approach to interrogate major modes of protein regulation. We generated datasets describing changes in protein and mRNA levels, protein subcellular localization, detergent insolubility, and post-translational modifications (PTMs) in the aging brain of killifish (Figure 2A and S3A). First, we captured proteome-wide variation in subcellular localization using an approach based on differential centrifugation coupled with quantitative mass spectrometry (LOPIT-DC) (*18*) and analyzed pools of adult (12 weeks post-hatching = wph) and old (39 wph) killifish brains (Figure S3B, Table S2). We used a list of well-annotated organelle markers (*88*) to evaluate organelle separation by LOPIT-DC (Figure S5A and S3C, D) and to confirm the reproducibility of organelles sedimentation between adult and old brains (Figure S3E). We then employed a tailored statistical approach (see methods, Figure S3F) to identify age-dependent changes in protein sedimentation profiles (Figure S5B, Table S2). The most prominent changes affected multiple mitochondrial and lysosomal proteins among others, including the mitochondrial transporters SLC25A32 and SLC25A18, and the lysosomal and vesicle trafficking proteins RAB14 and CCZ1 (Figure S5C). We interpret these alterations of sedimentation as an indication of partial reorganization of the mitochondrial and lysosomal proteome during aging that correlates with the well-described dysfunction of these organelles during aging and neurodegenerative diseases.

In parallel, we used the same pools of samples to assess age-dependent changes in protein solubility. We complemented our previous analysis of SDS insoluble aggregates in the killifish aging brain (*6*) with a more fine-grained analysis of protein solubility. Thus, we exposed brain homogenates to a series of detergent combinations of increasing strength, separated soluble and insoluble fractions by ultracentrifugation (as described in (*17*), Figure S2A, Table S2), and quantified protein abundances across fractions using mass spectrometry. Principal component analysis showed reproducible detergent-based fractionation in adult and old brains (Figure S2B) and GO enrichment analysis confirmed the expected partitioning of cellular components as a function of detergent strength (Figure S2C and S2D). In agreement with previous findings from other species (*11*, *89*) and the spontaneous age-related accumulation of protein aggregates in killifish brain (*5*–*7*), we observed an overall increase of protein detergent-insolubility in old samples (Figure S2E). By comparing detergent insolubility profiles between adult and old brains (Figure S2F-G), we identified 410 protein groups changing detergent insolubility with aging (Figure S5D, Table S2). While many of these proteins exhibited increased insolubility to detergents in old brains, there were instances where aging was linked to decreased insolubility to detergents. This indicates that factors other than protein aggregation, such as alterations in protein interactions or localization, could be responsible for the observed changes in detergent insolubility.

Next, we examined the effects of brain aging on multiple PTMs, using a sequential enrichment strategy followed by quantification of age-dependent changes in protein ubiquitylation, acetylation, and phosphorylation in the aging brain (Figure S4A, Table S3). We quantified PTM-carrying peptides normalized for protein changes (see methods, Figure S4) and identified age-related changes for 534 phosphorylated, 618 ubiquitylated, and 190 acetylated peptides (*P*<0.05, Figure S5E). The general increase in the number of affected PTM peptides with aging emphasized its overall impact on the proteome beyond protein abundance (Figure S5E-F). Integration of phosphorylation data with experimentally derived kinase-substrate relationships (*76*) indicates a remodeling of kinase signaling in the aging brain. Besides an increased activity (i.e., increased phosphorylation of predicted targets) for kinases involved in the regulation of immune responses, we reported enhanced activity for kinases of the protein kinase C family, e.g., PKN1, PKN2, PKCA, whose hyperactivation is linked to Alzheimer’s disease (*90*,*91*,*92*). Our data also reveals the decreased activity of kinases responsible for the phosphorylation of splicing factors and other RNA processing proteins, e.g. CDC2-like kinases 2 and 4 (CLK2 and CLK4, Figure S5G-H). These data suggest a convergence between aging and neurodegeneration concerning altered signaling pathways in the brain and hints at dysfunctional RNA processing in the aging brain.

To more systematically investigate the convergence between brain aging and neurodegenerative diseases, we queried our datasets for killifish orthologs of proteins encoded by genes that have been genetically linked to neurodegeneration in humans (Table S4). We found several of these proteins to be affected by aging in killifish in at least one of the proteomic datasets analyzed (Figure S5I). These include changes in subcellular fractionation and detergent insolubility (Figure S6A-B), as well as 23 PTM sites conserved between killifish and humans (Figure S6C-D-E). The microtubule-associated protein Tau (MAPT) was notably affected by aging across multiple proteomic layers. MAPT showed a prominent increase in detergent insolubility in old brains (Figure S5D), an alteration associated with human aging and neurodegenerative diseases (*93*–*95*). Additionally, we detected an age-dependent increase in phosphorylation and ubiquitylation of conserved residues in the microtubule-binding domain (MBD) of MAPT, a region sensitive to PTMs and associated with Tau pathological aggregation (Figure S5J and S6D) (*96*), (*95*, *97*). We validated the spontaneous increase of MAPT/Tau phosphorylation in old killifish brains using immunofluorescence staining for a conserved phosphorylated epitope of Tau (AT100) (Figure S5K).

Together, our analyses comprehensively establish how aging affects the brain proteome along multiple axes beyond protein abundance, using a consistent model organism and age groups. This thorough characterization of the proteome reveals several potential connections between aging, specific molecular events, and genetic factors associated with neurodegeneration, which are relevant to humans. To make this resource easily accessible to the scientific community, we have developed a web application at xxxxxx

---

**Figure S1:**
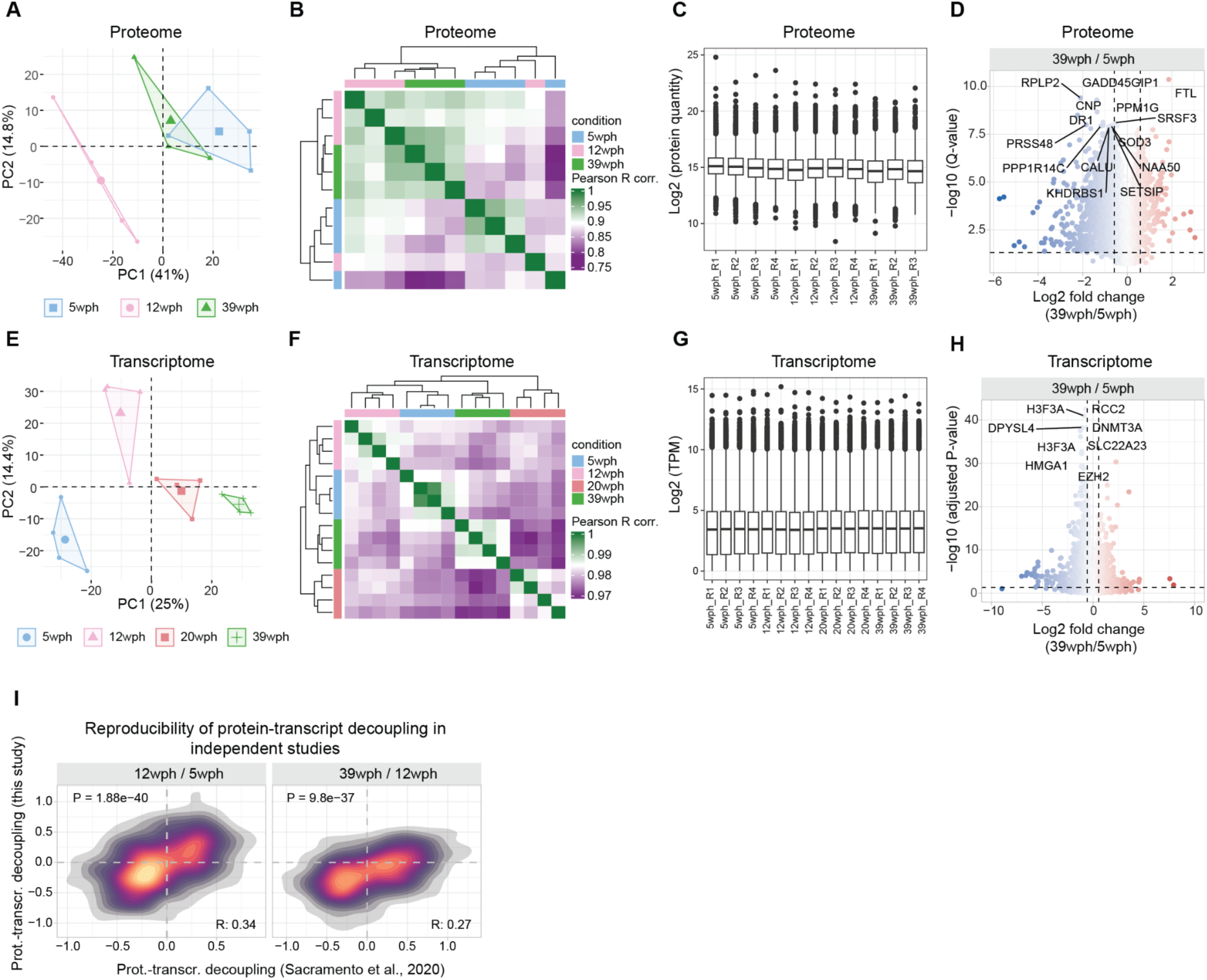
Proteome and transcriptome characterization of the killifish aging brain. A) Principal component analysis of proteomics data. B) Correlation heatmap between samples from the aging brain proteome data. Pairwise Pearson’s R correlation coefficient was calculated on the log2 transformed protein abundances. C) Boxplot displaying the distribution of log2 transformed and normalized protein abundances. D) Volcano plot highlighting significant protein abundance changes in the aging brain (39 wph vs. 5 wph). Dashed lines indicate the threshold used to select differentially abundant proteins (absolute log2 FC > 0.58 and -log10 Q-value < 0.05) E) Principal component analysis of transcriptomics data. F) Correlation heatmap between samples from the aging brain transcriptome data. Pairwise Pearson’s R correlation coefficient was calculated on the log2 transformed transcript per million reads (TPM). G) Boxplot displaying the distribution of log2 transformed and normalized transcript counts (TPM). H) Volcano plot highlighting significant transcript abundance changes in the aging brain (39 wph vs 5 wph). Dashed lines indicate the threshold used to select differentially expressed genes (absolute log2 FC > 0.58 and -log10 Adjusted P-value < 0.05). For displaying purposes, the X-axis range was limited to a -10:10 range leading to the exclusion of 1 gene. I) 2-D density plot showing the correlation between protein-transcript decoupling during aging in this study, displayed on the y-axis, and protein-transcript decoupling described in (*6*) (x-axis). Related to Figure 1 and Table S1.

**Figure S2:**
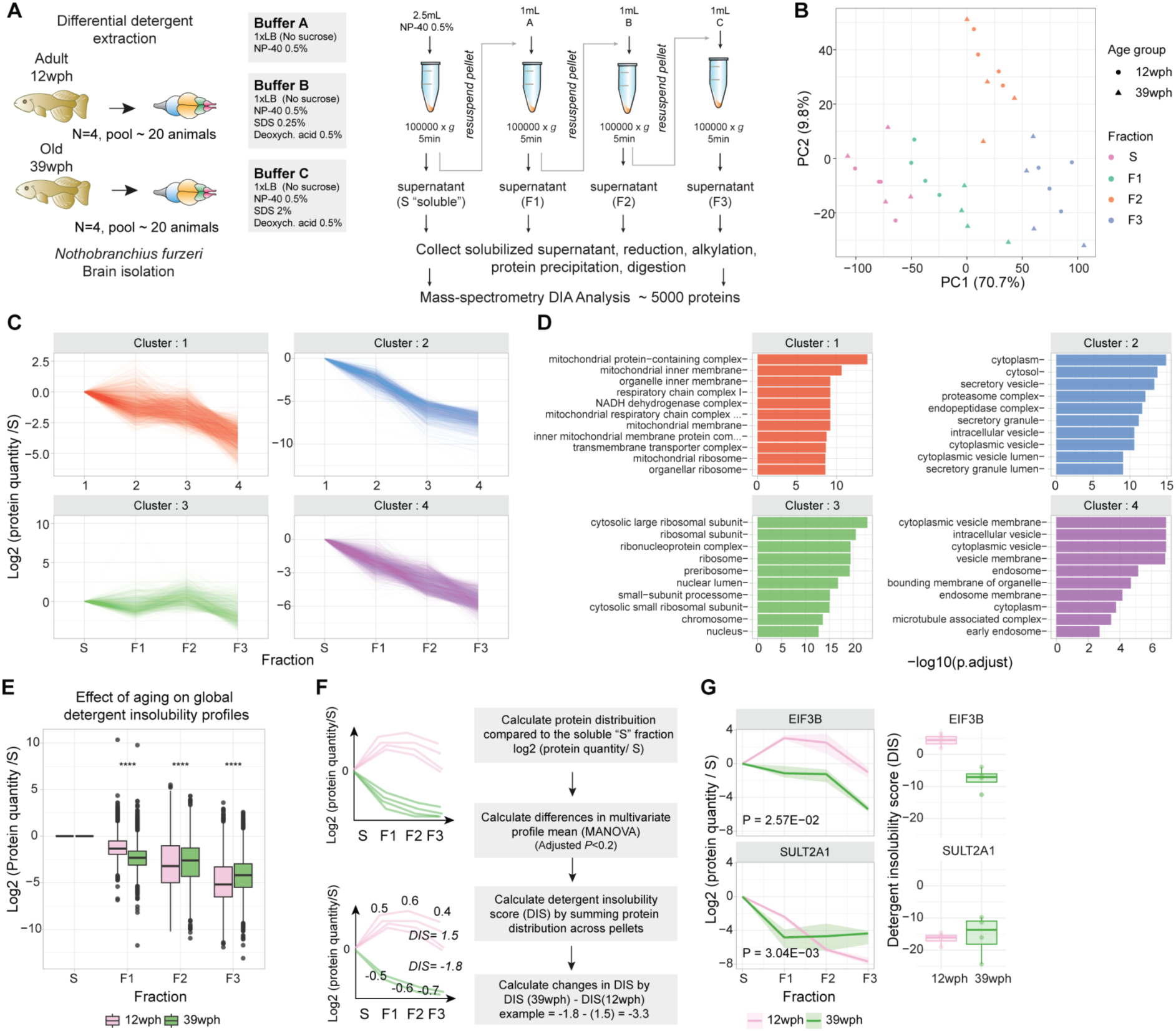
Protein detergent insolubility changes in the killifish aging brain. A) Scheme of the differential detergent extraction experiment. The protocol was adapted to brain tissue from ((*17*) see methods). B) Principal component analysis based on proteomics data from fractions obtained by differential detergent extraction. C) K-means clustering of detergent insolubility profiles. On the y-axis, the log2 protein quantity relative to the soluble “S” fraction, each profile represents the median across both conditions and (N=4 pools) replicates. D) GO enrichment overrepresentation analysis (ORA) of proteins assigned to each cluster against the rest of the identified proteome. On the x-axis, the -log10 of the adjusted P-value (Holm correction) of the Fisher’s Test is reported. Colors refer to the different clusters displayed in panel C. E) Boxplot depicting detergent insolubility profiles for all the proteins quantified across age groups. The y-axis indicates the log2 transformed value of protein quantity in each fraction relative to the soluble (S) fraction. Asterisks indicate the results of a two-sample Wilcoxon test. F) Computational strategy used for calculating differences in detergent insolubility profile across age groups. A MANOVA test was performed on each protein profile to detect significant changes in the multivariate mean between 12 wph (adult) and 39 wph (old samples), N=4 pools per age group. The detergent insolubility score (DIS) was calculated by summing the log2 protein quantity (relative to the soluble S fraction). Higher DIS indicate proteins that are relatively more abundant in insoluble fractions (F1:F3) than the soluble one (S). G) Example profiles of top hits proteins displaying changes in detergent insolubility with aging. EIF3B is an example of a protein that displays decreased detergent insolubility with age, while SULT2A1 displays increased detergent insolubility with age. For the left panel, the y-axis represents the log2 protein quantity in each fraction relative to the first soluble (S) fraction. Dark lines indicate the median between replicates, while shaded areas represent 50% of the replicate distribution, N=4 pools per age group. On the right panel, boxplots show the Detergent insolubility score (calculated as the sum of the log2 protein quantity relative to the first soluble (S) fraction) for the same proteins. Related to Figure 2, S5 and Table S2. *P ≤ 0.05; **P ≤ 0.01, ***P ≤ 0.001, ****P ≤ 0.0001.

**Figure S3:**
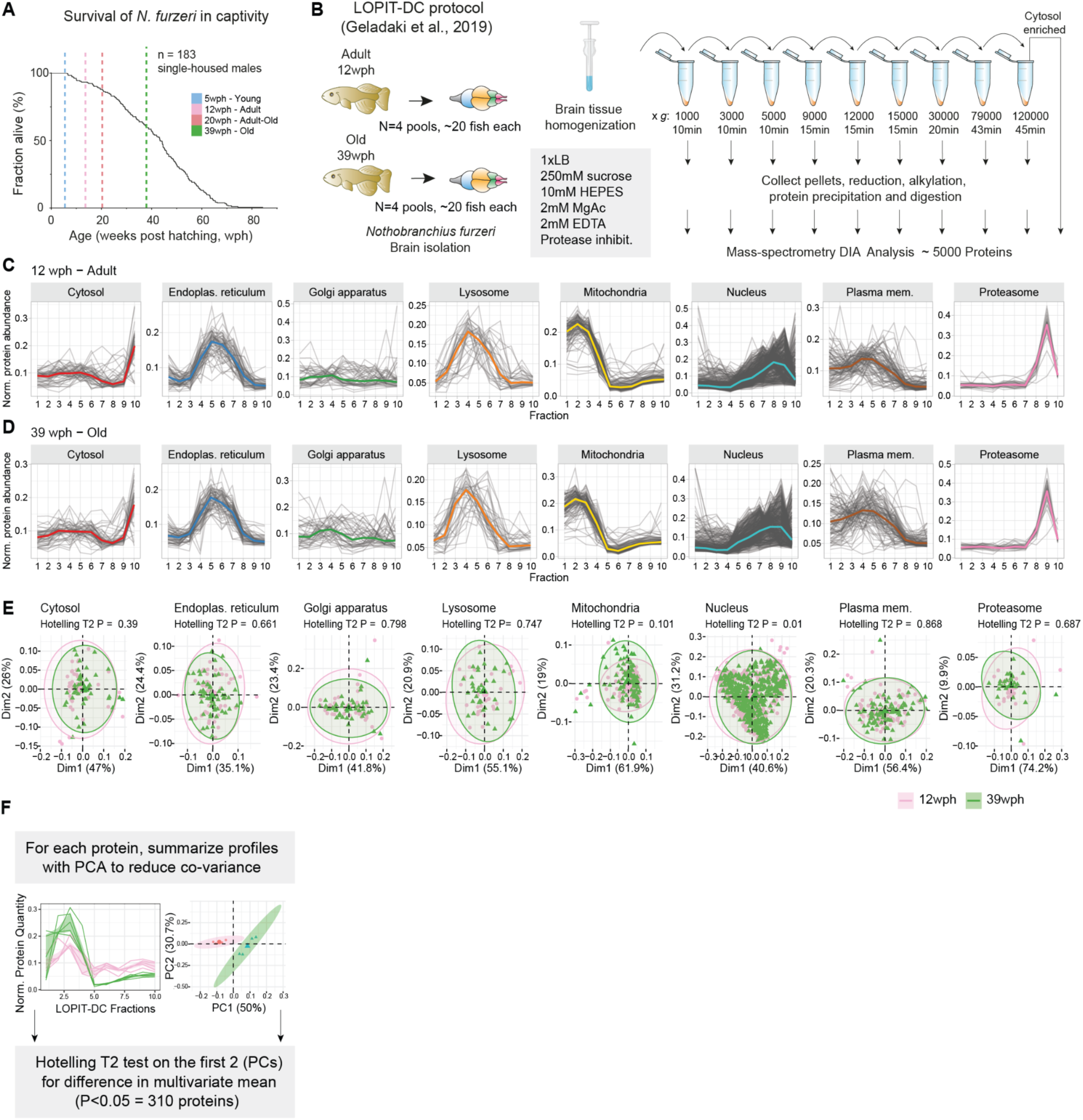
Subcellular fractionation of the killifish aging brain by LOPIT-DC. A) Survival curves of *Nothobranchius furzeri* MZM-0410 strain in captivity (data from (*98*)). The survival of *Nothobranchius furzeri* was investigated by tracking the occurrence of deaths starting at the age of 5 weeks post-hatching (wph), which corresponds to sexual maturity. This study includes data from four age groups highlighted by vertical dashed lines. The analyzed strain was derived from the wild with a median lifespan of 7-8 months. B) Scheme of the LOPIT-DC experiment. The protocol was adapted to brain tissue from (*18*) see methods for details. C-D) Organelle markers protein profiles from LOPIT-DC. The x-axis indicates the different fractions. The y-axis indicates protein abundance estimates derived from label-free Data Independent Acquisition mass spectrometry. Protein quantities were normalized by dividing the protein quantity in each fraction by the sum of the protein quantity along fractions. Each profile represents the median across replicates (N=4 pools). The median profiles of each organelle are highlighted by a colored solid line. Profiles obtained from adult (12 wph, panel C) and old (39 wph, panel D) fish are shown. E) Principal component analysis for different organelles markers in the LOPIT-DC fractions. Organelle markers from 12 wph (pink) and 39 wph (green) are shown. Each dot represents the median profile across (N=4 pools) replicate for each condition. F) Computational strategy used to identify age-related changes in protein sedimentation profiles. Related to Figure 2, Figure S5 and Table S2.

**Figure S4:**
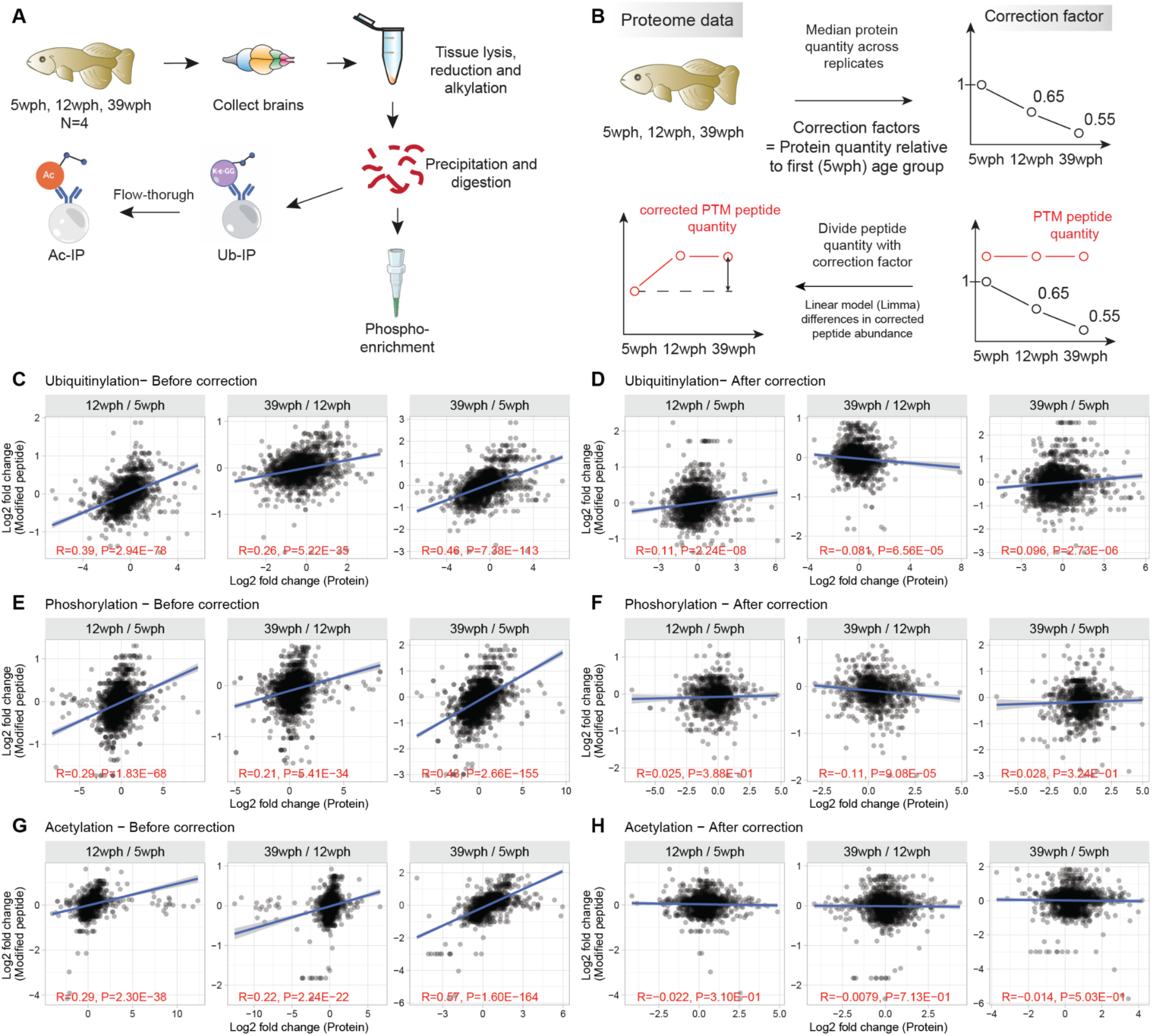
Analysis of protein post-translational modifications in the killifish aging brain. A) Workflow for the enrichment of post-translational modified peptides from in killifish brain. B) Correction strategy for detecting stoichiometric changes in post-translationally modified peptides. Correction factors were computed for each protein and condition relative to the 5 wph (young) age group. Quantities of the modified peptides were divided by the corresponding protein correction factor, and age-related changes were tested using *limma* (*75*). C-H) Relationship between age-related abundance changes of modified peptides vs. corresponding protein, before (left panels) and after (right panels) correction. The red text indicates the test results for the association between paired samples using Pearson’s product-moment correlation coefficients. Related to Figure 2, Figure S5 and Table S3.

**Figure S5:**
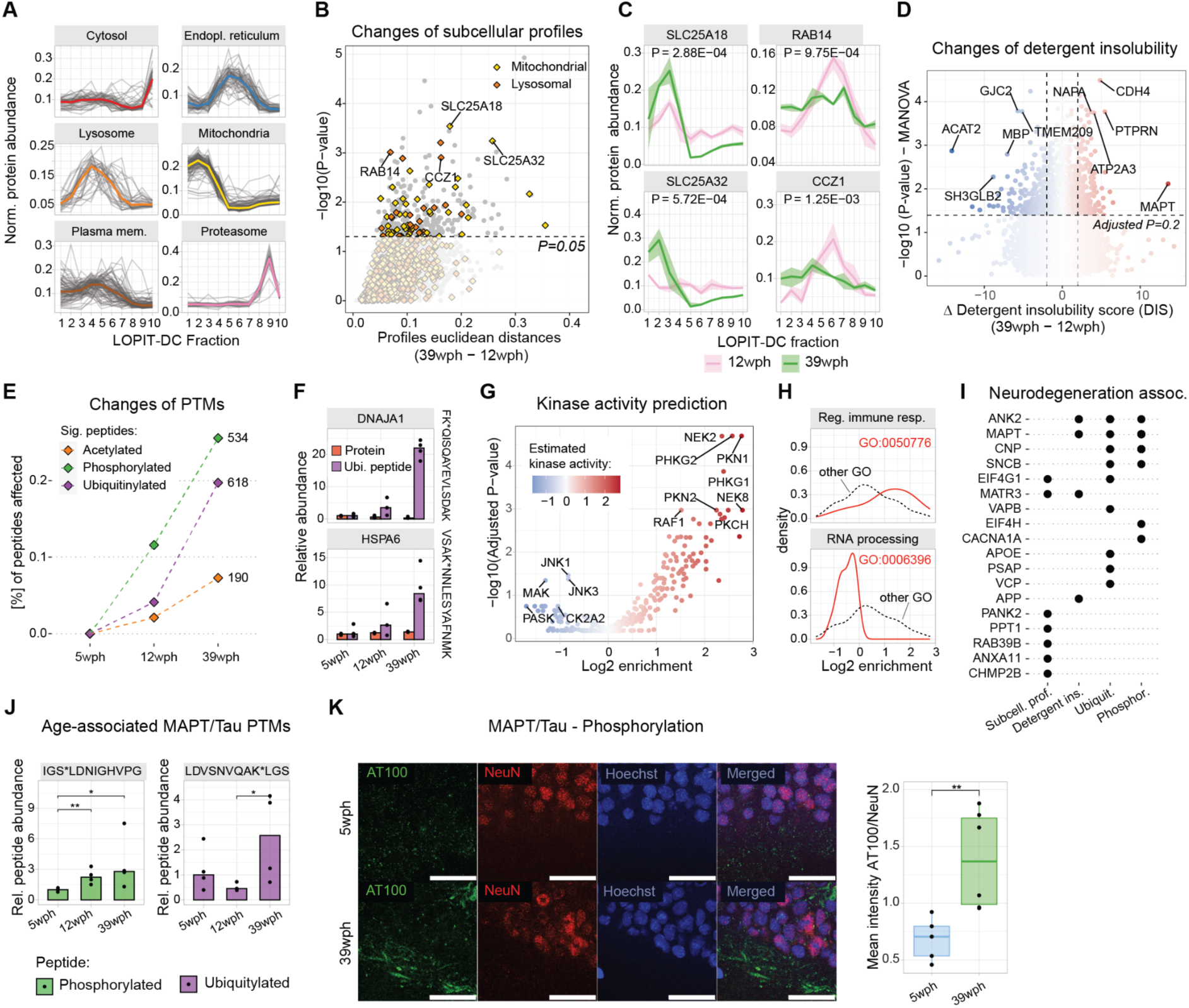
Aging affects protein subcellular localization, detergent insolubility and PTMs. A) Organelle markers protein profiles from LOPIT-DC (12 wph). The x-axis indicates the different fractions of the LOPIT-DC experiment. The y-axis indicates protein distribution across fractions. The median profiles of each organelle are highlighted by a colored solid line. B) Scatterplot depicting protein relocalization scores in the aging killifish brain. The x-axis indicates the median replicate Euclidean distance of the profiles between the two conditions. Y-axis indicates the -log10 P-value of the Hotelling T-squared test, between adult and old profiles (N=4 pools per age group). C) Examples of sedimentation profiles for selected proteins with altered subcellular fractionation profiles. In each of the plots, the x-axis indicates the 10 fractions obtained from LOPIT-DC, the y-axis indicates the total protein distribution along the 10 fractions for adult (pink) and old (green) fish. Shaded areas indicate 50% of the replicate distribution. P-values indicate the results of the Hotelling T2 test, (N=4 pools per age group). D) Volcano plot depicting protein detergent insolubility changes in the aging killifish brain. The x-axis indicates the difference in detergent insolubility score (see methods) expressed as old vs. adult. Higher values indicate increased detergent insolubility in the old brain. Y-axis indicates the -log10 of the MANOVA test between adult and old profiles (N=4 pools per age group). Significant changes are highlighted by dashed lines (MANOVA adjusted P<0.2 and absolute Δ Detergent insolubility score >2). E) Post-translationally modified peptides affected by aging. The y-axis (left) indicates the percentage of affected sites in each dataset when compared to the young samples (*P*<0.05, moderated Bayes T-test, N=3-4). F) Barplots showing relative abundances of ubiquitylated peptides from DNAJA1 and HSPA6 across age groups (purple bars). The corresponding protein abundances are displayed as reference (red bars), N=3-4. G) Volcano plot showing changes in estimated kinase activity (using the algorithm from (*76*)) based on phosphoproteomics data from old vs. young fish brains. The x-axis indicates changes in estimated kinase activity, the y-axis indicates FDR corrected -log10(P-value, Fisher’s test). H) Density distribution for kinases involved in the regulation of immune response (GO:0050776, upper panel) and RNA processing (GO:0006396, lower panel) against all other kinases from panel H. x-axis indicates the log2 Kinase activity enrichment value. I) Heatmap showing alterations of proteins linked to neurodegenerative diseases. Significant alterations in each dataset (P<0.05) are marked by black dots. J) Barplots displaying significantly changing (*P*<0.05, moderated Bayes T-test) MAPT/Tau phosphorylated (green) and ubiquitinated (purple) peptide. The values represent relative abundances to the young age group, after correction for protein changes (see methods, Figure S4B). Asterisks indicate the P-value of the moderated Bayes T-test (N=3-4). K) (Left panel) Immunofluorescence stainings for phosphorylated (AT100) Tau in brain cryo-sections of young and old N. furzeri. The stainings were normalized over the amount of NeuN in order to account for the different amounts of neuronal cells between young and old (N=5) animals. Scale bars = 20μm. (Right panel) Boxplot representation of mean intensity for phosphorylated Tau normalized over the amount of NeuN. P-value indicates the results of a two-sample Wilcoxon test. *P ≤ 0.05; **P ≤ 0.01, ***P ≤ 0.001, ****P ≤ 0.0001. Related to Table S2,S3,S4.

**Figure S6:**
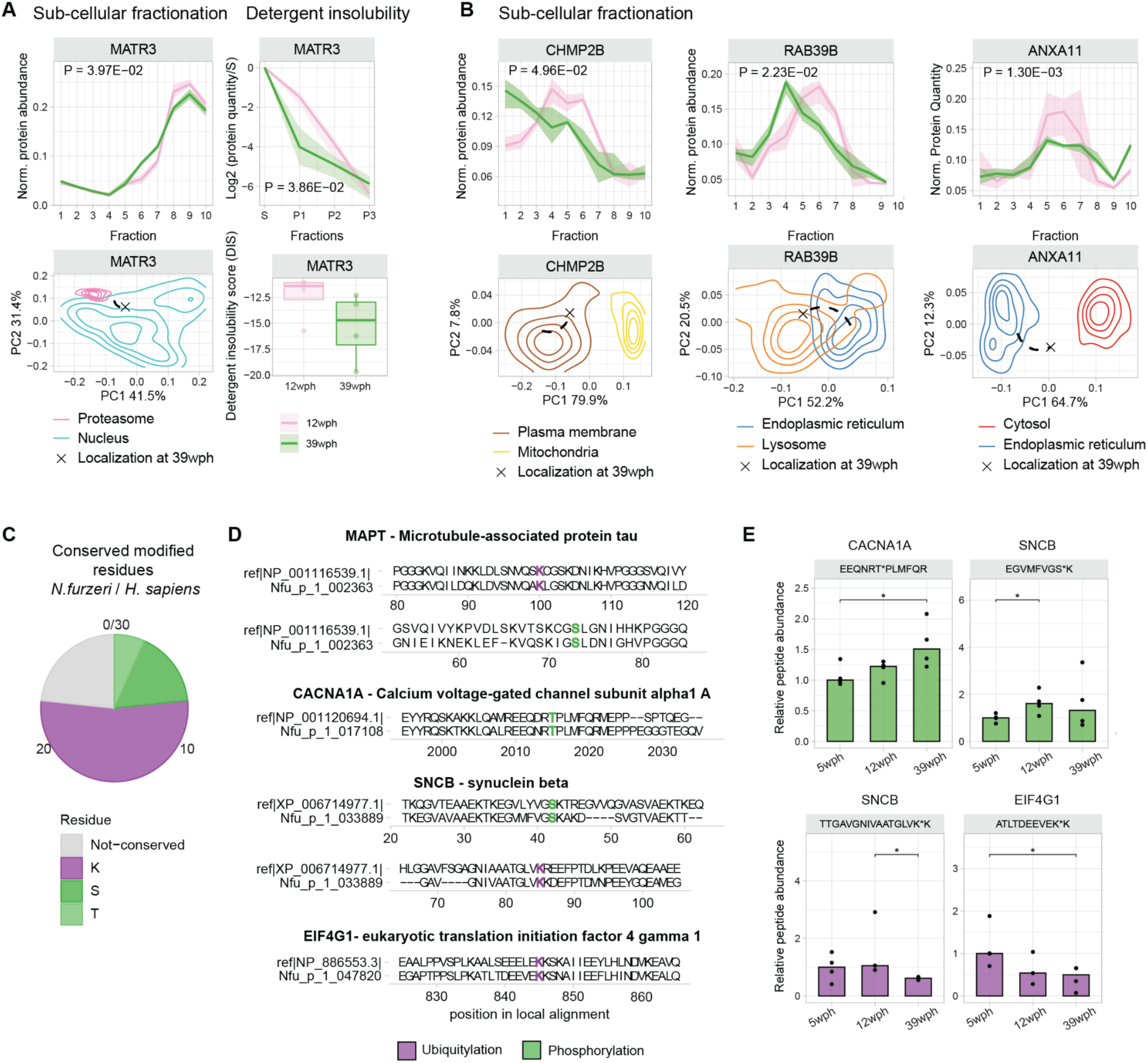
Age-associated alterations of proteins linked to human neurodegenerative disorders. A-B) Examples of proteins changing their subcellular localization profile or detergent insolubility. The top panels indicate either subcellular fractionation profiles (as in Figure 1D) or detergent insolubility profiles. For subcellular fractionation, in each of the plots, the x-axis indicates the 10 fractions obtained from LOPIT-DC and the y-axis indicates the total protein distribution along the 10 fractions for adult (12 wph, pink) and old (39 wph, green) fish. Shaded areas indicate 50% of the (N=4 pools) replicate distribution. P-values indicate the results of the Hotelling T2 test. For detergent insolubility profiles, the x-axis indicates the different detergent insolubility fractions: S=soluble, F1:F3=fractions after solubilization with buffers of increasing detergent strength (see methods, Figure S2A). The y-axis indicates log2 protein quantities relative to the soluble (S) fraction. The shaded area indicates 50% of the distribution across N=4 pools per age group. In the bottom panels, the PCA plot represents relocalization for each protein. The contour line represents the density distribution of the different organelles (calculated as the median between 12 wph and 39 wph), and the position of the protein at 39 wph is highlighted with a cross. The organelles represented are the ones that possess the higher absolute changes in the log2 ratios between Euclidean distances from the protein in the two age groups. Only for panel A, the boxplot on the right side indicates the detergent insolubility score in the two age groups. C) Pieplot showing conserved modified residues between *Nothobranchius furzeri* and humans that display changes in abundance with aging. Data refers to proteins involved in neurodegenerative diseases in humans. D) Local sequence alignments between *Nothobranchius furzeri* proteins (bottom sequence) and best human BLAST hit (upper sequence) for different proteins involved in neurodegenerative diseases. Modified residues are highlighted in purple (ubiquitylation) and green (phosphorylation). E) Barplots displaying significantly changing (*P*<0.05, moderated Bayes T-test) of modified peptides for the proteins shown in panel D. Asterisks indicate the P-value of the moderated Bayes T-test (N=3-4). The values represent relative abundances to the young (5 wph) age group after correction for protein changes (see methods, Figure S4B). Related to Figure S5 and Table S4.

**Figure S7:**
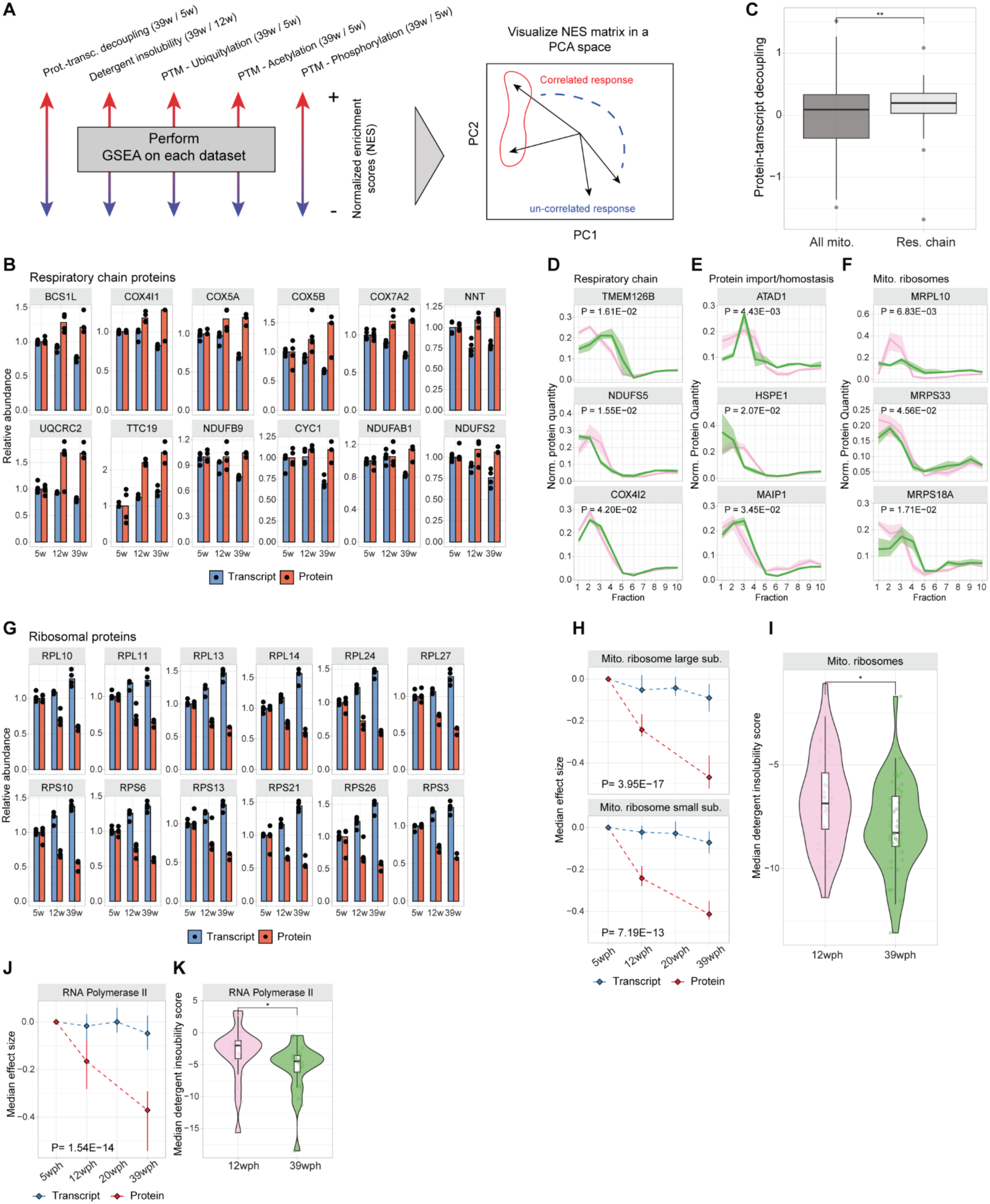
Alterations of ribosomal and respiratory chain proteins. A) Scheme of data integration strategy. For each dataset, a gene set enrichment analysis (GSEA) was performed using GO terms for cellular components. The normalized enrichment scores (NES) from each dataset were combined in a matrix and used as input for principal component analysis. B) Barplot showing transcript and protein abundances for oxidative phosphorylation protein. All the values were normalized to the 5 wph (young) age group (set to 1), N=3-4. C) Boxplot depicting the distribution of protein-transcript decoupling values (as defined in Figure 2A) for oxidative phosphorylation (light gray) proteins against the rest of the mitochondrial proteome (dark gray). Asterisks indicate the results of a two-sample Wilcoxon test. D-F) Examples of mitochondrial proteins that display changes in subcellular fractionation with aging. The x-axis indicates the 10 fractions obtained from LOPIT-DC, and the y-axis indicates the total protein distribution along the 10 fractions for adult (12 wph, pink) and old (39 wph, green) animals. Shaded areas indicate 50% of the replicate distribution from N=4 pools per group. P-values indicate the results of the Hotelling T2 test. G) Barplot showing transcript and protein abundances for cytoplasmic ribosomal protein. All the values were normalized to the 5 wph (young) age group (set to 1), N=3-4. H) Line plot showing the trajectories for transcriptome (blue) and proteome (red) of mitochondrial large and small ribosomal subunits. Each point summarizes the median distribution of the log2 ratio of the quantities relative to the first (5 wph) age group, while the line bars indicate 50% of the distributions. P-values indicate the results of a MANOVA test run on the two multivariate distributions, N=3-4. I) Violin plot displaying detergent insolubility score for proteins of the mitochondrial ribosome (GO:0005761). Each dot represents the median insolubility score of each protein across N=4 pools per age group; asterisks indicate the results of a two-sample Wilcoxon test. J) Line plot showing the trajectories for transcriptome (blue) and proteome (red) for RNA Polymerase II enzyme. Each point summarizes the median distribution of the log2 ratio of the quantities relative to the first (5 wph) age group, while the line bars indicate 50% of the distributions. P-values indicate the results of a MANOVA test run on the two multivariate distributions, N=3-4. K) Violin plot displaying detergent insolubility score for proteins of the RNA Polymerase II enzyme (GO:0016591). Each dot represents the median insolubility score of each protein across N=4 pools per age group; asterisks indicate the results of a two-sample Wilcoxon test. *P ≤ 0.05; **P ≤ 0.01, ***P ≤ 0.001, ****P ≤ 0.0001. Related to Figure 3.

**Figure S8:**
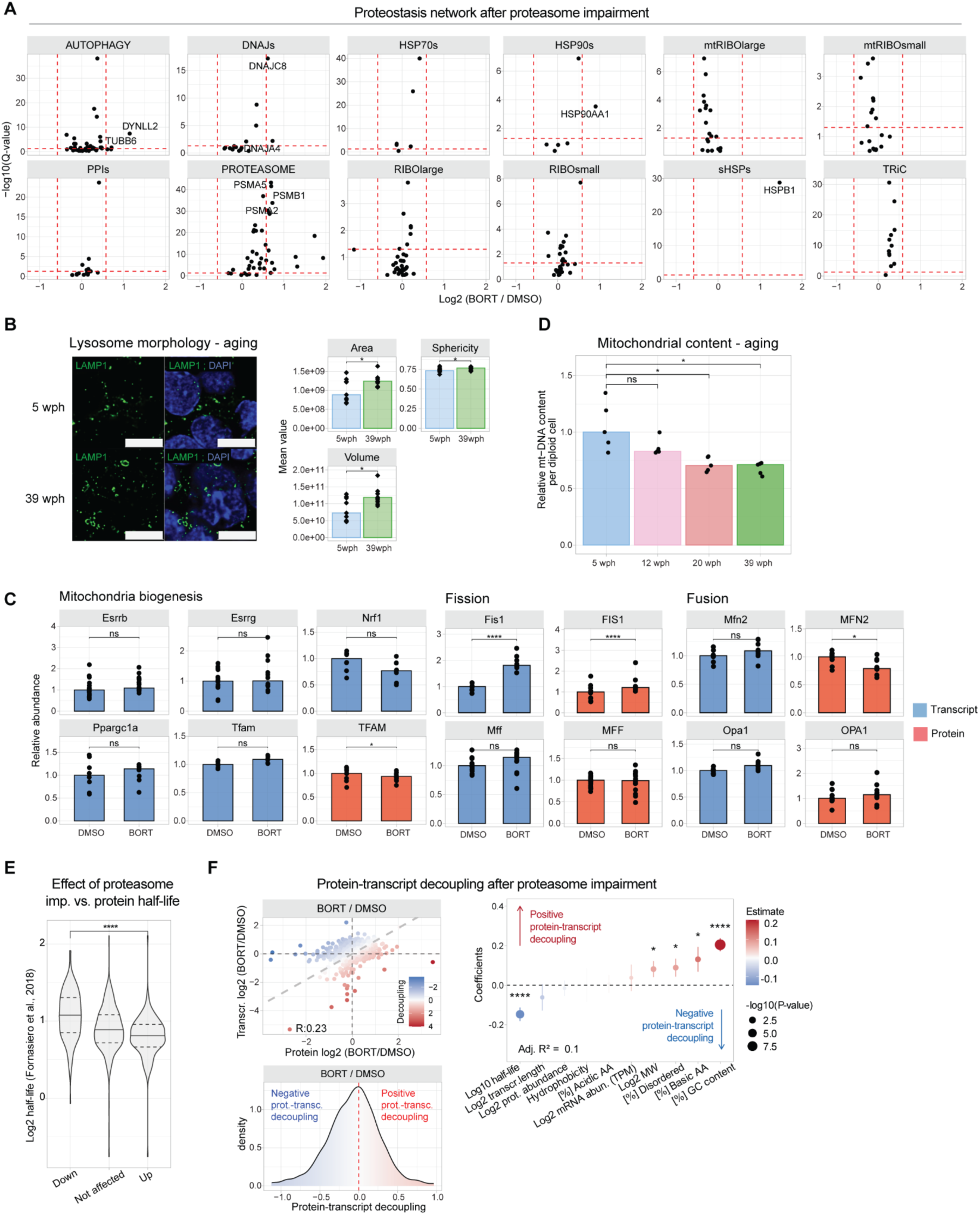
Effect of proteasome impairment on the killifish brain. A) Protein abundance changes induced by proteasome impairment for different components of the proteostasis network. B) (Right panel) Immunofluorescence stainings for lysosome (LAMP1) in brain cryo-sections of young (light blue) and old (green) *Nothobranchius furzeri*. Scale bars = 5μm. (Left panel) Barplot representation of lysosome morphology features in young (light blue) and old (green) samples. The y-axis represents the mean value of the different morphology features in each of the replicates (N=6).C) Effect of proteasome impairment on mitochondrial transcripts and proteins. For protein data, asterisks indicate the Q-value of the differential abundance testing performed with a two-sample T-test on the peptide abundances. For transcript data, asterisks indicate the Adjusted P-value of the differential abundance testing. N=10. D) Quantification of mitochondrial DNA (mt-DNA) from killifish brains during aging. Relative mtDNA copy number was calculated using real-time quantitative PCR with primers for 16S rRNA mitochondrial gene and Cdkn2a/b nuclear gene for normalization (N=5). Asterisks indicate the results of two-sample Wilcoxon tests. E) Violin plot showing the distribution of up and down-regulated proteins in response to proteasome impairment against their half-life as quantified in (*16*). Asterisks indicate the results of a two-samples Wilcoxon test. F) (Top left panel) Scatterplot comparing protein- (x-axis) and transcript-level (y-axis) fold changes in killifish after treatment with bortezomib. The color of each dot represents the decoupling score calculated as the difference between log2 transformed fold changes measured at the protein and transcript levels. Grey dashed lines indicate the equal changes between transcript and protein and, therefore, a zero decoupling score. (Bottom left panel) Density distribution of decoupling scores for comparing bortezomib vs. DMSO. On the right part, highlighted in red, are protein “gain” events (increase in protein abundance compared to the transcript), while on the left, in blue, are protein “loss” events (decrease in protein abundance compared to the transcript). (Right panel) Multiple linear regression analysis of decoupling scores in response to proteasome impairment based on biophysical features of transcripts or proteins as predictors. The x-axis indicates the estimate of the regression coefficient for each feature, while the size of the dots and asterisks represent the -log10 P-values of the F-test. *P ≤ 0.05; **P ≤ 0.01, ***P ≤ 0.001, ****P ≤ 0.0001. Related to Figure 3 and Table S5.

**Figure S9:**
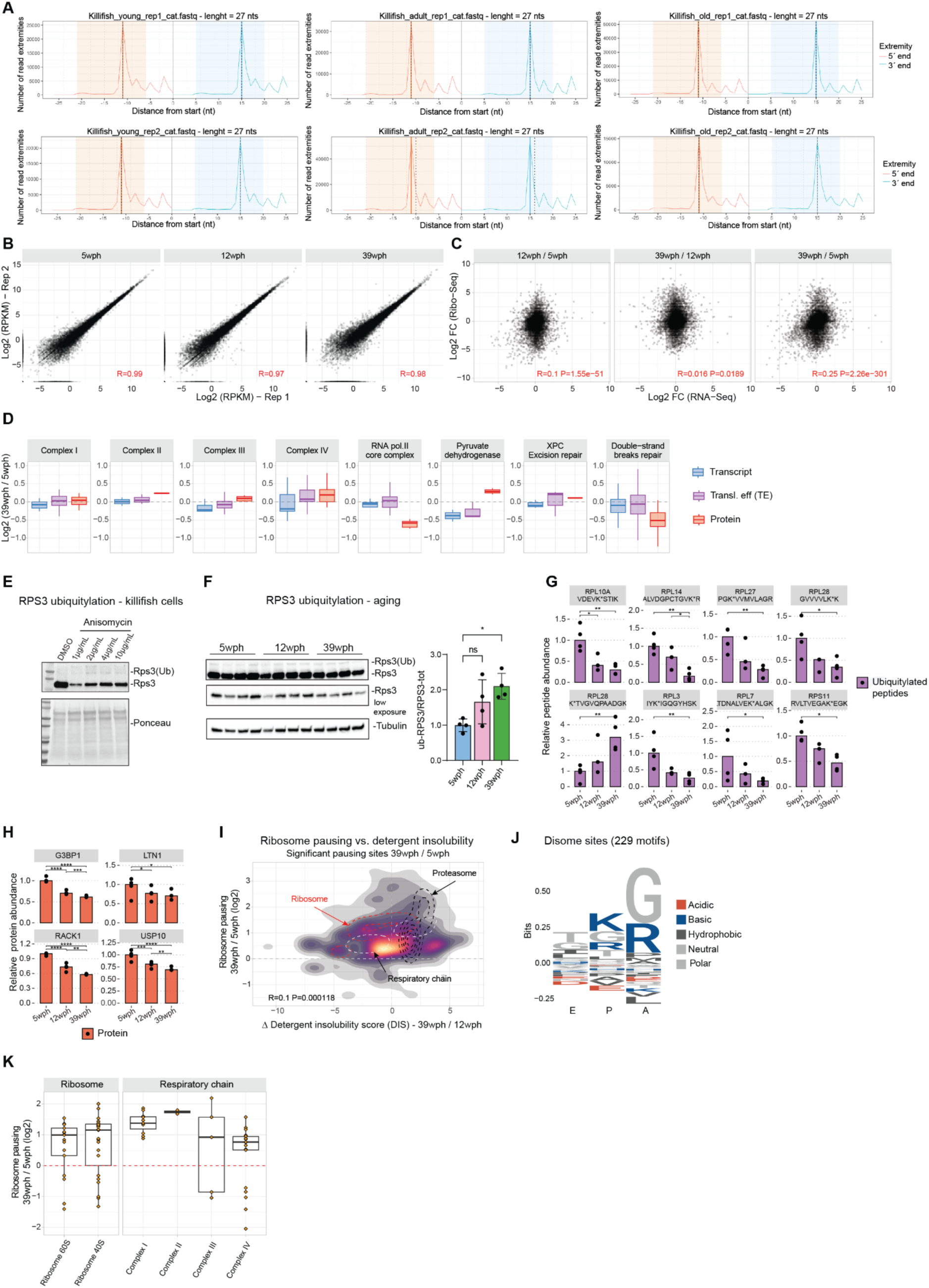
Ribosome profiling in the killifish aging brain. A) Tri-nucleotide plot showing characteristic triplet periodicity. The x-axis represents the distance from the starting codon (in nucleotide) and the y-axis the number of reads. B) Scatterplot showing the correlation between replicates for the Ribo-Seq experiment. On the different axis, the log2(RPKM) values from the different replicates are shown. C) Scatterplot showing the correlation between log2 fold changes for ribosome occupancy (y-axis) and changes in the transcriptome (x-axis) for different aging steps. D) Boxplot displaying differential modes of regulation for different protein complexes. On the x-axis are displayed the different datasets: Transcriptome (green), Translation efficiency (purple), and Proteome (red). E) Immunoblot to detect RPS3 ubiquitylation in killifish cells treated with Anisomycin, which inhibits translation elongation and causes ribotoxic stress (*99*) for 24 hours. F) Immunoblot to detect RPS3 ubiquitylation across age groups. Barplot shows the ratio between the total RPS3 and its ubiquitylated fraction during aging. Asterisks indicate the results of an ordinary one-way ANOVA test (N=4). G) Barplots displaying significantly changing (*P*<0.05, moderated Bayes T-test) of ubiquitin-modified peptides for ribosomal proteins. Asterisks indicate the P-value of the moderated Bayes T-test (N=3-4). The values represent relative abundances to the young (5 wph) age group after correction for protein changes (see methods, Figure S4B). H) Barplot showing normalized protein abundance (relative to the first, 5 wph, age group set to 1) for factors associated with Ribosome-Quality-Control (RQC) pathways. The y-axis represents protein abundances relative to the first (5 wph) age groups. Asterisks indicate the Q-value of the differential abundance testing performed with a two-sample T-test on the peptide abundances, N=3,4 pools per group. I) 2-D density plot showing the relation between significant changes in pausing (Adjusted P-value < 0.05) displayed on the y-axis and changes in detergent insolubility metrics (x-axis). Each point in the distribution represents a significantly altered pausing site. Contour lines indicate the distribution of cytoplasmic ribosomes (red), Proteasome (black), and oxidative phosphorylation (white). J) Weblogo for disome pausing sites that display a strong increase in pausing (Pause score > 10). The y-axis displays the relative frequencies of the different residues, while the x-axis displays the different ribosome positions (E, P, A). K) Boxplot showing the distributions of pausing sites for cytoplasmic ribosomes (left panel) and respiratory chain complexes (right). Each dot represents a significantly altered (Adjusted P-value < 0.05) pausing site. The Y axis represents the log2 fold changes in pausing between 39 wph and 5 wph. *P ≤ 0.05; **P ≤ 0.01, ***P ≤ 0.001, ****P ≤ 0.0001. Related to Figure 4 and Table S6.

## References

1. J. Labbadia, R. I. Morimoto, The biology of proteostasis in aging and disease. Annu. Rev. Biochem. (2015) (available at https://www.annualreviews.org/doi/abs/10.1146/annurev-biochem-060614-033955).

2. M. S. Hipp, P. Kasturi, F. U. Hartl, The proteostasis network and its decline in ageing. Nat. Rev. Mol. Cell Biol. 20, 421–435 (2019).

3. D. R. Valenzano, E. Terzibasi, A. Cattaneo, L. Domenici, A. Cellerino, Temperature affects longevity and age-related locomotor and cognitive decay in the short-lived fish Nothobranchius furzeri. Aging Cell. 5, 275–278 (2006).

4. S. Bagnoli, B. Fronte, C. Bibbiani, T. E. Terzibasi, A. Cellerino, Quantification of noradrenergic-, dopaminergic-, and tectal-neurons during aging in the short-lived killifish Nothobranchius furzeri. Aging Cell. 21 (2022), doi:10.1111/acel.13689.

5. H. Matsui, N. Kenmochi, K. Namikawa, Age- and α-Synuclein-Dependent Degeneration of Dopamine and Noradrenaline Neurons in the Annual Killifish Nothobranchius furzeri. Cell Rep. 26, 1727–1733.e6 (2019).

6. E. Kelmer Sacramento, J. M. Kirkpatrick, M. Mazzetto, M. Baumgart, A. Bartolome, S. Di Sanzo, C. Caterino, M. Sanguanini, N. Papaevgeniou, M. Lefaki, D. Childs, S. Bagnoli, E. Terzibasi Tozzini, D. Di Fraia, N. Romanov, P. H. Sudmant, W. Huber, N. Chondrogianni, M. Vendruscolo, A. Cellerino, A. Ori, Reduced proteasome activity in the aging brain results in ribosome stoichiometry loss and aggregation. Mol. Syst. Biol. 16, e9596 (2020).

7. I. Harel, Y. R. Chen, I. Ziv, P. P. Singh, P. N. Negredo, U. Goshtchevsky, W. Wang, G. Astre, E. Moses, A. McKay, B. E. Machado, K. Hebestreit, S. Yin, A. S. Alvarado, D. F. Jarosz, A. Brunet, Identification of protein aggregates in the aging vertebrate brain with prion-like and phase separation properties. bioRxiv (2022), p. 2022.02.26.482115.

8. A. Louka, S. Bagnoli, J. Rupert, B. Esapa, G. G. Tartaglia, A. Cellerino, A. Pastore, T. E. Terzibasi, New lessons on TDP-43 from old N. furzeri killifish. Aging Cell. 21 (2022), doi:10.1111/acel.13517.

9. G. E. Janssens, A. C. Meinema, J. González, J. C. Wolters, A. Schmidt, V. Guryev, R. Bischoff, E. C. Wit, L. M. Veenhoff, M. Heinemann, Protein biogenesis machinery is a driver of replicative aging in yeast. Elife. 4, e08527 (2015).

10. Y.-N. Wei, H.-Y. Hu, G.-C. Xie, N. Fu, Z.-B. Ning, R. Zeng, P. Khaitovich, Transcript and protein expression decoupling reveals RNA binding proteins and miRNAs as potential modulators of human aging. Genome Biol. 16, 41 (2015).

11. D. M. Walther, P. Kasturi, M. Zheng, S. Pinkert, G. Vecchi, P. Ciryam, R. I. Morimoto, C. M. Dobson, M. Vendruscolo, M. Mann, F. U. Hartl, Widespread Proteome Remodeling and Aggregation in Aging C. elegans. Cell. 161 (2015), doi:10.1016/j.cell.2015.03.032.

12. D. C. David, N. Ollikainen, J. C. Trinidad, M. P. Cary, A. L. Burlingame, C. Kenyon, Widespread Protein Aggregation as an Inherent Part of Aging in C. elegans. PLoS Biol. 8, e1000450 (2010).

13. Y. Takemon, J. M. Chick, I. Gerdes Gyuricza, D. A. Skelly, O. Devuyst, S. P. Gygi, G. A. Churchill, R. Korstanje, Proteomic and transcriptomic profiling reveal different aspects of aging in the kidney. Elife. 10 (2021), doi:10.7554/eLife.62585.

14. I. Gerdes Gyuricza, J. M. Chick, G. R. Keele, A. G. Deighan, S. C. Munger, R. Korstanje, S. P. Gygi, G. A. Churchill, Genome-wide transcript and protein analysis highlights the role of protein homeostasis in the aging mouse heart. Genome Res. 32, 838–852 (2022).

15. F. Dick, O. B. Tysnes, G. W. Alves, G. S. Nido, C. Tzoulis, Altered transcriptome-proteome coupling indicates aberrant proteostasis in Parkinson’s disease. iScience. 26 (2023), doi:10.1016/j.isci.2023.105925.

16. E. F. Fornasiero, S. Mandad, H. Wildhagen, M. Alevra, B. Rammner, S. Keihani, F. Opazo, I. Urban, T. Ischebeck, M. S. Sakib, M. K. Fard, K. Kirli, T. P. Centeno, R. O. Vidal, R.-U. Rahman, E. Benito, A. Fischer, S. Dennerlein, P. Rehling, I. Feussner, S. Bonn, M. Simons, H. Urlaub, S. O. Rizzoli, Precisely measured protein lifetimes in the mouse brain reveal differences across tissues and subcellular fractions. Nat. Commun. 9, 1–17 (2018).

17. A. T. N. Tebbenkamp, D. R. Borchelt, Protein Aggregate Characterization in Models of Neurodegenerative Disease. Neuroproteomics, 85–91 (2009).

18. A. Geladaki, N. K. Britovšek, L. M. Breckels, T. S. Smith, O. L. Vennard, C. M. Mulvey, O. M. Crook, L. Gatto, K. S. Lilley, Combining LOPIT with differential ultracentrifugation for high-resolution spatial proteomics. Nature Communications. 10 (2019),, doi:10.1038/s41467-018-08191-w.

19. D. A. Gray, M. Tsirigotis, J. Woulfe, Ubiquitin, proteasomes, and the aging brain. Sci. Aging Knowledge Environ. 2003 (2003), doi:10.1126/sageke.2003.34.re6.

20. M. E. G. de Araujo, G. Liebscher, M. W. Hess, L. A. Huber, Lysosomal size matters. Traffic. 21, 60–75 (2020).

21. M. Stagi, Z. A. Klein, T. J. Gould, J. Bewersdorf, S. M. Strittmatter, Lysosome size, motility and stress response regulated by fronto-temporal dementia modifier TMEM106B. Mol. Cell. Neurosci. 61, 226–240 (2014).

22. C. M. Pickart, Back to the future with ubiquitin. Cell. 116, 181–190 (2004).

23. A. Ori, B. H. Toyama, M. S. Harris, T. Bock, M. Iskar, P. Bork, N. T. Ingolia, M. W. Hetzer, M. Beck, Integrated Transcriptome and Proteome Analyses Reveal Organ-Specific Proteome Deterioration in Old Rats. Cell Syst. 1, 224–237 (2015).

24. K. C. Stein, F. Morales-Polanco, J. van der Lienden, T. K. Rainbolt, J. Frydman, Ageing exacerbates ribosome pausing to disrupt cotranslational proteostasis. Nature. 601, 637–642 (2022).

25. C. Meyer, A. Garzia, P. Morozov, H. Molina, T. Tuschl, The G3BP1-Family-USP10 Deubiquitinase Complex Rescues Ubiquitinated 40S Subunits of Ribosomes Stalled in Translation from Lysosomal Degradation. Mol. Cell. 77, 1193–1205.e5 (2020).

26. R. Higgins, J. M. Gendron, L. Rising, R. Mak, K. Webb, S. E. Kaiser, N. Zuzow, P. Riviere, B. Yang, E. Fenech, X. Tang, S. A. Lindsay, J. C. Christianson, R. Y. Hampton, S. A. Wasserman, E. J. Bennett, The Unfolded Protein Response Triggers Site-Specific Regulatory Ubiquitylation of 40S Ribosomal Proteins. Mol. Cell. 59, 35–49 (2015).

27. L. L. Yan, C. L. Simms, F. McLoughlin, R. D. Vierstra, H. S. Zaher, Oxidation and alkylation stresses activate ribosome-quality control. Nat. Commun. 10, 5611 (2019).

28. A. K. Sharma, J. Venezian, A. Shiber, G. Kramer, B. Bukau, E. P. O’Brien, Combinations of slow-translating codon clusters can increase mRNA half-life in. Proc. Natl. Acad. Sci. U. S. A. 118 (2021), doi:10.1073/pnas.2026362118.

29. L. Y. Chan, C. F. Mugler, S. Heinrich, P. Vallotton, K. Weis, Non-invasive measurement of mRNA decay reveals translation initiation as the major determinant of mRNA stability. Elife. 7 (2018), doi:10.7554/eLife.32536.

30. D. C. Schwartz, R. Parker, mRNA decapping in yeast requires dissociation of the cap binding protein, eukaryotic translation initiation factor 4E. Mol. Cell. Biol. 20, 7933–7942 (2000).

31. D. Gaidatzis, L. Burger, M. Florescu, M. B. Stadler, Erratum: Analysis of intronic and exonic reads in RNA-seq data characterizes transcriptional and post-transcriptional regulation. Nat. Biotechnol. 34, 210 (2016).

32. E. W. Mills, R. Green, Ribosomopathies: There’s strength in numbers. Science. 358 (2017), doi:10.1126/science.aan2755.

33. R. K. Khajuria, M. Munschauer, J. C. Ulirsch, C. Fiorini, L. S. Ludwig, S. K. McFarland, N. J. Abdulhay, H. Specht, H. Keshishian, D. R. Mani, M. Jovanovic, S. R. Ellis, C. P. Fulco, J. M. Engreitz, S. Schütz, J. Lian, K. W. Gripp, O. K. Weinberg, G. S. Pinkus, L. Gehrke, A. Regev, E. S. Lander, H. T. Gazda, W. Y. Lee, V. G. Panse, S. A. Carr, V. G. Sankaran, Ribosome Levels Selectively Regulate Translation and Lineage Commitment in Human Hematopoiesis. Cell. 173, 90–103.e19 (2018).

34. W. L. Noderer, R. J. Flockhart, A. Bhaduri, A. J. Diaz de Arce, J. Zhang, P. A. Khavari, C. L. Wang, Quantitative analysis of mammalian translation initiation sites by FACS-seq. Mol. Syst. Biol. 10, 748 (2014).

35. C. López-Otín, M. A. Blasco, L. Partridge, M. Serrano, G. Kroemer, Hallmarks of aging: An expanding universe. Cell. 186, 243–278 (2023).

36. M. Deschênes, B. Chabot, The emerging role of alternative splicing in senescence and aging. Aging Cell. 16, 918–933 (2017).

37. A. Gyenis, J. Chang, J. J. P. G. Demmers, S. T. Bruens, S. Barnhoorn, R. M. C. Brandt, M. P. Baar, M. Raseta, K. W. J. Derks, J. H. J. Hoeijmakers, J. Pothof, Genome-wide RNA polymerase stalling shapes the transcriptome during aging. Nat. Genet. 55, 268–279 (2023).

38. B. Schumacher, J. Pothof, J. Vijg, J. H. J. Hoeijmakers, The central role of DNA damage in the ageing process. Nature. 592, 695–703 (2021).

39. M. Bhadra, P. Howell, S. Dutta, C. Heintz, W. B. Mair, Alternative splicing in aging and longevity. Hum. Genet. 139, 357–369 (2020).

40. M. Bozukova, C. Nikopoulou, N. Kleinenkuhnen, D. Grbavac, K. Goetsch, P. Tessarz, Aging is associated with increased chromatin accessibility and reduced polymerase pausing in liver. Mol. Syst. Biol. 18, e11002 (2022).

41. C. Debès, A. Papadakis, S. Grönke, Ö. Karalay, L. S. Tain, A. Mizi, S. Nakamura, O. Hahn, C. Weigelt, N. Josipovic, A. Zirkel, I. Brusius, K. Sofiadis, M. Lamprousi, Y.-X. Lu, W. Huang, R. Esmaillie, T. Kubacki, M. R. Späth, B. Schermer, T. Benzing, R.-U. Müller, A. Antebi, L. Partridge, A. Papantonis, A. Beyer, Ageing-associated changes in transcriptional elongation influence longevity. Nature. 616, 814–821 (2023).

42. P. Kapahi, J. Vijg, Aging--lost in translation? N. Engl. J. Med. 361, 2669–2670 (2009).

43. Y. Gonskikh, N. Polacek, Alterations of the translation apparatus during aging and stress response. Mech. Ageing Dev. 168, 30–36 (2017).

44. T. Ingram, L. Chakrabarti, Proteomic profiling of mitochondria: what does it tell us about the ageing brain? Aging. 8, 3161–3179 (2016).

45. J. C. Heiby, A. Ori, Organelle dysfunction and its contribution to metabolic impairments in aging and age-related diseases. Current Opinion in Systems Biology. 30, 100416 (2022).

46. N. Miyoshi, H. Oubrahim, P. B. Chock, E. R. Stadtman, Age-dependent cell death and the role of ATP in hydrogen peroxide-induced apoptosis and necrosis. Proc. Natl. Acad. Sci. U. S. A. 103, 1727–1731 (2006).

47. B. P. Braeckman, K. Houthoofd, J. R. Vanfleteren, Aging Cell, in press.

48. D. Gkotsi, R. Begum, T. Salt, G. Lascaratos, C. Hogg, K.-Y. Chau, A. H. V. Schapira, G. Jeffery, Recharging mitochondrial batteries in old eyes. Near infra-red increases ATP. Exp. Eye Res. 122, 50–53 (2014).

49. L. Espada, A. Dakhovnik, P. Chaudhari, A. Martirosyan, L. Miek, T. Poliezhaieva, Y. Schaub, A. Nair, N. Döring, N. Rahnis, O. Werz, A. Koeberle, J. Kirkpatrick, A. Ori, M. A. Ermolaeva, Loss of metabolic plasticity underlies metformin toxicity in aged Caenorhabditis elegans. Nat Metab. 2, 1316–1331 (2020).

50. A. A. Bazzini, F. Del Viso, M. A. Moreno-Mateos, T. G. Johnstone, C. E. Vejnar, Y. Qin, J. Yao, M. K. Khokha, A. J. Giraldez, Codon identity regulates mRNA stability and translation efficiency during the maternal-to-zygotic transition. EMBO J. 35, 2087–2103 (2016).

51. P. T. da Silva, Y. Zhang, E. Theodorakis, L. D. Martens, V. A. Yépez, V. Pelechano, J. Gagneur, Cellular energy regulates mRNA translation and degradation in a codon-specific manner. bioRxiv (2023), p. 2023.04.06.535836.

52. M. Ximerakis, S. L. Lipnick, B. T. Innes, S. K. Simmons, X. Adiconis, D. Dionne, B. A. Mayweather, L. Nguyen, Z. Niziolek, C. Ozek, V. L. Butty, R. Isserlin, S. M. Buchanan, S. S. Levine, A. Regev, G. D. Bader, J. Z. Levin, L. L. Rubin, Single-cell transcriptomic profiling of the aging mouse brain. Nat. Neurosci. 22 (2019), doi:10.1038/s41593-019-0491-3.

53. A. Tyshkovskiy, S. Ma, A. V. Shindyapina, S. Tikhonov, S.-G. Lee, P. Bozaykut, J. P. Castro, A. Seluanov, N. J. Schork, V. Gorbunova, S. E. Dmitriev, R. A. Miller, V. N. Gladyshev, Distinct longevity mechanisms across and within species and their association with aging. Cell. 186, 2929–2949.e20 (2023).

54. Q. Yu, H. Xiao, M. P. Jedrychowski, D. K. Schweppe, J. Navarrete-Perea, J. Knott, J. Rogers, E. T. Chouchani, S. P. Gygi, Sample multiplexing for targeted pathway proteomics in aging mice. Proc. Natl. Acad. Sci. U. S. A. 117, 9723–9732 (2020).

55. S. Koyuncu, R. Loureiro, H. J. Lee, P. Wagle, M. Krueger, D. Vilchez, Rewiring of the ubiquitinated proteome determines ageing in C. elegans. Nature. 596, 285–290 (2021).

56. V. Kluever, B. Russo, S. Mandad, N. H. Kumar, M. Alevra, A. Ori, S. O. Rizzoli, H. Urlaub, A. Schneider, E. F. Fornasiero, Protein lifetimes in aged brains reveal a proteostatic adaptation linking physiological aging to neurodegeneration. Science Advances. 8, eabn4437 (2022).

57. Z. Wu, I. Tantray, J. Lim, S. Chen, Y. Li, Z. Davis, C. Sitron, J. Dong, S. Gispert, G. Auburger, O. Brandman, X. Bi, M. Snyder, B. Lu, MISTERMINATE Mechanistically Links Mitochondrial Dysfunction with Proteostasis Failure. Mol. Cell. 75 (2019), doi:10.1016/j.molcel.2019.06.031.

58. S. Rimal, Y. Li, R. Vartak, J. Geng, I. Tantray, S. Li, S. Huh, H. Vogel, C. Glabe, L. T. Grinberg, S. Spina, W. W. Seeley, S. Guo, B. Lu, Inefficient quality control of ribosome stalling during APP synthesis generates CAT-tailed species that precipitate hallmarks of Alzheimer’s disease. Acta neuropathologica communications. 9 (2021), doi:10.1186/s40478-021-01268-6.

59. S. Li, Z. Wu, I. Tantray, Y. Li, S. Chen, J. Dong, S. Glynn, H. Vogel, M. Snyder, B. Lu, Quality-control mechanisms targeting translationally stalled and C-terminally extended poly(GR) associated with ALS/FTD. Proc. Natl. Acad. Sci. U. S. A. 117 (2020), doi:10.1073/pnas.2005506117.

60. R. Aviner, T.-T. Lee, V. B. Masto, D. Gestaut, K. H. Li, R. Andino, J. Frydman, Ribotoxic collisions on CAG expansions disrupt proteostasis and stress responses in Huntington’s Disease. bioRxiv (2022), p. 2022.05.04.490528.

61. Y. Aman, T. Schmauck-Medina, M. Hansen, R. I. Morimoto, A. K. Simon, I. Bjedov, K. Palikaras, A. Simonsen, T. Johansen, N. Tavernarakis, D. C. Rubinsztein, L. Partridge, G. Kroemer, J. Labbadia, E. F. Fang, Autophagy in healthy aging and disease. Nature Aging. 1, 634–650 (2021).

62. S. Safaiyan, N. Kannaiyan, N. Snaidero, S. Brioschi, K. Biber, S. Yona, A. L. Edinger, S. Jung, M. J. Rossner, M. Simons, Age-related myelin degradation burdens the clearance function of microglia during aging. Nat. Neurosci. 19, 995–998 (2016).

63. T. Stoeger, R. A. Grant, A. C. McQuattie-Pimentel, K. R. Anekalla, S. S. Liu, H. Tejedor-Navarro, B. D. Singer, H. Abdala-Valencia, M. Schwake, M.-P. Tetreault, H. Perlman, W. E. Balch, N. S. Chandel, K. M. Ridge, J. I. Sznajder, R. I. Morimoto, A. V. Misharin, G. R. S. Budinger, L. A. Nunes Amaral, Aging is associated with a systemic length-associated transcriptome imbalance. Nature Aging. 2, 1191–1206 (2022).

64. K. Buczak, J. M. Kirkpatrick, F. Truckenmueller, D. Santinha, L. Ferreira, S. Roessler, S. Singer, M. Beck, A. Ori, Spatially resolved analysis of FFPE tissue proteomes by quantitative mass spectrometry. Nat. Protoc. 15, 2956–2979 (2020).

65. S. Di Sanzo, K. Spengler, A. Leheis, J. M. Kirkpatrick, T. L. Rändler, T. Baldensperger, T. Dau, C. Henning, L. Parca, C. Marx, Z.-Q. Wang, M. A. Glomb, A. Ori, R. Heller, Mapping protein carboxymethylation sites provides insights into their role in proteostasis and cell proliferation. Nat. Commun. 12, 6743 (2021).

66. D. R. Bentley, S. Balasubramanian, H. P. Swerdlow, G. P. Smith, J. Milton, C. G. Brown, K. P. Hall, D. J. Evers, C. L. Barnes, H. R. Bignell, J. M. Boutell, J. Bryant, R. J. Carter, R. Keira Cheetham, A. J. Cox, D. J. Ellis, M. R. Flatbush, N. A. Gormley, S. J. Humphray, L. J. Irving, M. S. Karbelashvili, S. M. Kirk, H. Li, X. Liu, K. S. Maisinger, L. J. Murray, B. Obradovic, T. Ost, M. L. Parkinson, M. R. Pratt, I. M. J. Rasolonjatovo, M. T. Reed, R. Rigatti, C. Rodighiero, M. T. Ross, A. Sabot, S. V. Sankar, A. Scally, G. P. Schroth, M. E. Smith, V. P. Smith, A. Spiridou, P. E. Torrance, S. S. Tzonev, E. H. Vermaas, K. Walter, X. Wu, L. Zhang, M. D. Alam, C. Anastasi, I. C. Aniebo, D. M. D. Bailey, I. R. Bancarz, S. Banerjee, S. G. Barbour, P. A. Baybayan, V. A. Benoit, K. F. Benson, C. Bevis, P. J. Black, A. Boodhun, J. S. Brennan, J. A. Bridgham, R. C. Brown, A. A. Brown, D. H. Buermann, A. A. Bundu, J. C. Burrows, N. P. Carter, N. Castillo, M. Chiara E. Catenazzi, S. Chang, R. Neil Cooley, N. R. Crake, O. O. Dada, K. D. Diakoumakos, B. Dominguez-Fernandez, D. J. Earnshaw, U. C. Egbujor, D. W. Elmore, S. S. Etchin, M. R. Ewan, M. Fedurco, L. J. Fraser, K. V. Fuentes Fajardo, W. Scott Furey, D. George, K. J. Gietzen, C. P. Goddard, G. S. Golda, P. A. Granieri, D. E. Green, D. L. Gustafson, N. F. Hansen, K. Harnish, C. D. Haudenschild, N. I. Heyer, M. M. Hims, J. T. Ho, A. M. Horgan, K. Hoschler, S. Hurwitz, D. V. Ivanov, M. Q. Johnson, T. James, T. A. Huw Jones, G.-D. Kang, T. H. Kerelska, A. D. Kersey, I. Khrebtukova, A. P. Kindwall, Z. Kingsbury, P. I. Kokko-Gonzales, A. Kumar, M. A. Laurent, C. T. Lawley, S. E. Lee, X. Lee, A. K. Liao, J. A. Loch, M. Lok, S. Luo, R. M. Mammen, J. W. Martin, P. G. McCauley, P. McNitt, P. Mehta, K. W. Moon, J. W. Mullens, T. Newington, Z. Ning, B. Ling Ng, S. M. Novo, M. J. O’Neill, M. A. Osborne, A. Osnowski, O. Ostadan, L. L. Paraschos, L. Pickering, A. C. Pike, A. C. Pike, D. Chris Pinkard, D. P. Pliskin, J. Podhasky, V. J. Quijano, C. Raczy, V. H. Rae, S. R. Rawlings, A. Chiva Rodriguez, P. M. Roe, J. Rogers, M. C. Rogert Bacigalupo, N. Romanov, A. Romieu, R. K. Roth, N. J. Rourke, S. T. Ruediger, E. Rusman, R. M. Sanches-Kuiper, M. R. Schenker, J. M. Seoane, R. J. Shaw, M. K. Shiver, S. W. Short, N. L. Sizto, J. P. Sluis, M. A. Smith, J. Ernest Sohna Sohna, E. J. Spence, K. Stevens, N. Sutton, L. Szajkowski, C. L. Tregidgo, G. Turcatti, S. vandeVondele, Y. Verhovsky, S. M. Virk, S. Wakelin, G. C. Walcott, J. Wang, G. J. Worsley, J. Yan, L. Yau, M. Zuerlein, J. Rogers, J. C. Mullikin, M. E. Hurles, N. J. McCooke, J. S. West, F. L. Oaks, P. L. Lundberg, D. Klenerman, R. Durbin, A. J. Smith, Accurate whole human genome sequencing using reversible terminator chemistry. Nature. 456, 53–59 (2008).

67. A. Dobin, C. A. Davis, F. Schlesinger, J. Drenkow, C. Zaleski, S. Jha, P. Batut, M. Chaisson, T. R. Gingeras, STAR: ultrafast universal RNA-seq aligner. Bioinformatics. 29, 15–21 (2012).

68. T. Smith, A. Heger, I. Sudbery, UMI-tools: modeling sequencing errors in Unique Molecular Identifiers to improve quantification accuracy. Genome Res. 27, 491–499 (2017).

69. Y. Liao, G. K. Smyth, W. Shi, featureCounts: an efficient general purpose program for assigning sequence reads to genomic features. Bioinformatics. 30, 923–930 (2013).

70. M. I. Love, W. Huber, S. Anders, Moderated estimation of fold change and dispersion for RNA-seq data with DESeq2. Genome Biol. 15, 550 (2014).

71. N. J. McGlincy, N. T. Ingolia, Transcriptome-wide measurement of translation by ribosome profiling. Methods. 126, 112–129 (2017).

72. S. Bagnoli, E. Terzibasi Tozzini, A. Cellerino, Immunofluorescence and Aggresome Staining of Nothobranchius furzeri Cryosections. Cold Spring Harb. Protoc. (2023), doi:10.1101/pdb.prot107791.

73. O. M. Crook, L. M. Breckels, K. S. Lilley, P. D. W. Kirk, L. Gatto, A Bioconductor workflow for the Bayesian analysis of spatial proteomics. F1000Res. 8, 446 (2019).

74. L. M. Breckels, C. M. Mulvey, K. S. Lilley, L. Gatto, A Bioconductor workflow for processing and analysing spatial proteomics data. F1000Res. 5, 2926 (2016).

75. M. E. Ritchie, B. Phipson, D. Wu, Y. Hu, C. W. Law, W. Shi, G. K. Smyth, limma powers differential expression analyses for RNA-sequencing and microarray studies. Nucleic Acids Res. 43, e47 (2015).

76. J. L. Johnson, T. M. Yaron, E. M. Huntsman, A. Kerelsky, J. Song, A. Regev, T. Y. Lin, K. Liberatore, D. M. Cizin, B. M. Cohen, N. Vasan, Y. Ma, K. Krismer, J. T. Robles, B. van de Kooij, A. E. van Vlimmeren, N. Andrée-Busch, N. F. Käufer, M. V. Dorovkov, A. G. Ryazanov, Y. Takagi, E. R. Kastenhuber, Goncalves, B. D. Hopkins, O. Elemento, D. J. Taatjes, A. Maucuer, A. Yamashita, A. Degterev, M. Uduman, J. Lu, S. D. Landry, B. Zhang, I. Cossentino, R. Linding, J. Blenis, P. V. Hornbeck, B. E. Turk, M. B. Yaffe, L. C. Cantley, An atlas of substrate specificities for the human serine/threonine kinome. Nature. 613 (2023), doi:10.1038/s41586-022-05575-3.

77. T. Wu, E. Hu, S. Xu, M. Chen, P. Guo, Z. Dai, T. Feng, L. Zhou, W. Tang, L. Zhan, X. Fu, S. Liu, X. Bo, G. Yu, clusterProfiler 4.0: A universal enrichment tool for interpreting omics data. Innovation (Camb). 2, 100141 (2021).

78. S. F. Altschul, W. Gish, W. Miller, E. W. Myers, D. J. Lipman, Basic local alignment search tool. J. Mol. Biol. 215, 403–410 (1990).

79. K. Strimmer, fdrtool: a versatile R package for estimating local and tail area-based false discovery rates. Bioinformatics. 24, 1461–1462 (2008).

80. F. Cunningham, J. E. Allen, J. Allen, J. Alvarez-Jarreta, M. R. Amode, I. M. Armean, O. Austine-Orimoloye, A. G. Azov, I. Barnes, R. Bennett, A. Berry, J. Bhai, A. Bignell, K. Billis, S. Boddu, L. Brooks, M. Charkhchi, C. Cummins, L. Da Rin Fioretto, C. Davidson, K. Dodiya, S. Donaldson, B. El Houdaigui, T. El Naboulsi, R. Fatima, C. G. Giron, T. Genez, J. G. Martinez, C. Guijarro-Clarke, A. Gymer, M. Hardy, Z. Hollis, T. Hourlier, T. Hunt, T. Juettemann, V. Kaikala, M. Kay, I. Lavidas, T. Le, D. Lemos, J. C. Marugán, S. Mohanan, A. Mushtaq, M. Naven, D. N. Ogeh, A. Parker, A. Parton, M. Perry, I. Piližota, I. Prosovetskaia, M. P. Sakthivel, A. I. A. Salam, B. M. Schmitt, H. Schuilenburg, D. Sheppard, J. G. Pérez-Silva, W. Stark, E. Steed, K. Sutinen, R. Sukumaran, D. Sumathipala, M.-M. Suner, M. Szpak, A. Thormann, F. F. Tricomi, D. Urbina-Gómez, A. Veidenberg, T. A. Walsh, B. Walts, N. Willhoft, A. Winterbottom, E. Wass, M. Chakiachvili, B. Flint, A. Frankish, S. Giorgetti, L. Haggerty, S. E. Hunt, G. R. IIsley, J. E. Loveland, F. J. Martin, B. Moore, J. M. Mudge, M. Muffato, E. Perry, M. Ruffier, J. Tate, D. Thybert, S. J. Trevanion, S. Dyer, P. W. Harrison, K. L. Howe, A. D. Yates, D. R. Zerbino, P. Flicek, Ensembl 2022. Nucleic Acids Res. 50, D988–D995 (2022).

81. A. R. Quinlan, I. M. Hall, BEDTools: a flexible suite of utilities for comparing genomic features. Bioinformatics. 26, 841–842 (2010).

82. D. J. Pagliarini, S. E. Calvo, B. Chang, S. A. Sheth, S. B. Vafai, S.-E. Ong, G. A. Walford, C. Sugiana, A. Boneh, W. K. Chen, D. E. Hill, M. Vidal, J. G. Evans, D. R. Thorburn, S. A. Carr, V. K. Mootha, A mitochondrial protein compendium elucidates complex I disease biology. Cell. 134, 112–123 (2008).

83. M. Martin, Cutadapt removes adapter sequences from high-throughput sequencing reads. EMBnet J. 17, 10 (2011).

84. B. Langmead, C. Trapnell, M. Pop, S. L. Salzberg, Ultrafast and memory-efficient alignment of short DNA sequences to the human genome. Genome Biol. 10, R25 (2009).

85. F. Lauria, T. Tebaldi, P. Bernabò, E. J. N. Groen, T. H. Gillingwater, G. Viero, riboWaltz: Optimization of ribosome P-site positioning in ribosome profiling data. PLoS Comput. Biol. 14, e1006169 (2018).

86. M. C. F. Thomsen, M. Nielsen, Seq2Logo: a method for construction and visualization of amino acid binding motifs and sequence profiles including sequence weighting, pseudo counts and two-sided representation of amino acid enrichment and depletion. Nucleic Acids Res. 40, W281–7 (2012).

87. G. H. Putri, S. Anders, P. T. Pyl, J. E. Pimanda, F. Zanini, Analysing high-throughput sequencing data in Python with HTSeq 2.0. Bioinformatics. 38, 2943–2945 (2022).

88. L. Gatto, L. M. Breckels, K. S. Lilley, Assessing sub-cellular resolution in spatial proteomics experiments. Curr. Opin. Chem. Biol. 48, 123–149 (2019).

89. G. Vecchi, P. Sormanni, B. Mannini, A. Vandelli, G. G. Tartaglia, C. M. Dobson, F. U. Hartl, M. Vendruscolo, Proteome-wide observation of the phenomenon of life on the edge of solubility. Proc. Natl. Acad. Sci. U. S. A. 117, 1015–1020 (2020).

90. S. I. Alfonso, J. A. Callender, B. Hooli, C. E. Antal, K. Mullin, M. A. Sherman, S. E. Lesné, M. Leitges, A. C. Newton, R. E. Tanzi, R. Malinow, Gain-of-function mutations in protein kinase Cα (PKCα) may promote synaptic defects in Alzheimer’s disease. Sci. Signal. 9 (2016), doi:10.1126/scisignal.aaf6209.

91. N. Morshed, M. J. Lee, F. H. Rodriguez, D. A. Lauffenburger, D. Mastroeni, F. M. White, Quantitative phosphoproteomics uncovers dysregulated kinase networks in Alzheimer’s disease. Nature Aging. 1, 550–565 (2021).

92. B. Bai, X. Wang, Y. Li, P.-C. Chen, K. Yu, K. K. Dey, J. M. Yarbro, X. Han, B. M. Lutz, S. Rao, Y. Jiao, J. M. Sifford, J. Han, M. Wang, H. Tan, T. I. Shaw, J.-H. Cho, S. Zhou, H. Wang, M. Niu, A. Mancieri, K. A. Messler, X. Sun, Z. Wu, V. Pagala, A. A. High, W. Bi, H. Zhang, H. Chi, V. Haroutunian, B. Zhang, T. G. Beach, G. Yu, J. Peng, Deep Multilayer Brain Proteomics Identifies Molecular Networks in Alzheimer’s Disease Progression. Neuron. 106, 700 (2020).

93. A. L. Guillozet, S. Weintraub, D. C. Mash, M. M. Mesulam, Neurofibrillary tangles, amyloid, and memory in aging and mild cognitive impairment. Arch. Neurol. 60, 729–736 (2003).

94. S. Chatterjee, M. Sealey, E. Ruiz, C. M. Pegasiou, K. Brookes, S. Green, A. Crisford, M. Duque-Vasquez, E. Luckett, R. Robertson, P. Richardson, G. Vajramani, P. Grundy, D. Bulters, C. Proud, M. Vargas-Caballero, A. Mudher, Age-related changes in tau and autophagy in human brain in the absence of neurodegeneration. PLoS One. 18, e0262792 (2023).

95. Y. Wang, E. Mandelkow, Tau in physiology and pathology. Nat. Rev. Neurosci. 17, 22–35 (2015).

96. L. Li, Y. Jiang, J.-Z. Wang, R. Liu, X. Wang, Tau Ubiquitination in Alzheimer’s Disease. Front. Neurol. 12 (2022), doi:10.3389/fneur.2021.786353.

97. D. Datta, S. N. Leslie, M. Wang, Y. M. Morozov, S. Yang, S. Mentone, C. Zeiss, A. Duque, P. Rakic, T. L. Horvath, C. H. van Dyck, A. C. Nairn, A. F. T. Arnsten, Age-related calcium dysregulation linked with tau pathology and impaired cognition in non-human primates. Alzheimers. Dement. 17, 920–932 (2021).

98. M. Baumgart, S. Priebe, M. Groth, N. Hartmann, U. Menzel, L. Pandolfini, P. Koch, M. Felder, M. Ristow, C. Englert, R. Guthke, M. Platzer, A. Cellerino, Longitudinal RNA-Seq Analysis of Vertebrate Aging Identifies Mitochondrial Complex I as a Small-Molecule-Sensitive Modifier of Lifespan. Cell Syst. 2, 122–132 (2016).

99. M. S. Iordanov, D. Pribnow, J. L. Magun, T. H. Dinh, J. A. Pearson, S. L. Chen, B. E. Magun, Ribotoxic stress response: activation of the stress-activated protein kinase JNK1 by inhibitors of the peptidyl transferase reaction and by sequence-specific RNA damage to the alpha-sarcin/ricin loop in the 28S rRNA. Mol. Cell. Biol. 17, 3373–3381 (1997).

